# A core set of neural states underlying naturalistic memory reactivation in the posterior medial cortex

**DOI:** 10.1101/2024.12.11.627957

**Authors:** Yoonjung Lee, Hongmi Lee, Janice Chen

**Affiliations:** Department of Psychological and Brain Sciences, Johns Hopkins University, Baltimore, MD 21218, USA; Department of Psychological Sciences, Purdue University, West Lafayette, IN 47907, USA

## Abstract

In the posterior midline default mode network, spatial activity patterns similar to those during the initial experience are reactivated during the successful recall of past events. Prior studies have shown that these event-specific activity patterns are consistent across individuals recalling a shared experience, suggesting that common functional responses underlying episodic recall do exist. However, the spatial organization of function during episodic encoding and subsequent recall, especially in the absence of external stimuli, remains poorly understood. To address this, we leverage fMRI data collected during the encoding and recall of naturalistic movies to identify a core set of neural states in the posterior medial cortex. These states are stimulus-locked, reactivated during recall, and have a shared spatial organization across brains (i.e., individuals). We show that a surprisingly small number of these states (16 states across hemispheres) is sufficient to achieve the same levels of reactivation in the posterior medial cortex as when using the standard methods of the field. Additionally, these states are linked to actions and social-affective features of events in the movies. Our findings elucidate the properties of a common, spatially organized code within the posterior default mode network which appears during natural recollection of memories.

## Introduction

While spatial organization of functional responses according to stimulus features is a common motif across much of sensory and mid-level mammalian cortex, little is known about how function is spatially organized during episodic recall of real-world events—a task which typically engages higher-order cortical regions. In low-level sensory processing areas, studies using fMRI have shown the presence of retinotopic maps in human early visual cortex^1–3^, and analogous tonotopic maps have been observed in early auditory cortex^4,5^. Further downstream in ventral temporal cortex, neural responses are selective to specific object categories, such as faces and objects^6,7^, with representational features such as animacy and real-world object size varying regularly across the cortical surface^8^ (see ref.^9^ for a review). However, such functional organization is typically assessed during the presence of external stimulation, such as when subjects are viewing gratings or images, or listening to sounds. In the default mode network (DMN)^10,11^, which occupies an apex position atop the brain’s cortical hierarchy, input from multiple sensory modalities is thought to be merged into more abstract, highly invariant representations^12^. DMN responses to naturalistic visual and auditory narrative stimuli are locked to semantic content and relatively insensitive to differences in input modality^13,14^ or language^15^. Furthermore, the DMN overlaps substantially with regions in which stimulus information is “reactivated” during retrieval of prior episodes^16,17^, even when no stimulus is available at the time of test. To what extent are these invariant, abstract, internally-driven representations in the DMN spatially organized, and how?

A number of recent studies provide evidence that spatially organized functional responses in DMN areas are likely to exist. Drawing on Event Segmentation Theory (EST)^18^, these studies model a continuous audiovisual movie as a series of discrete “events”, where “event boundaries” are defined by human judgments of when major perceptual or conceptual changes occur in the depicted scenario (e.g., moving to a new location; switching to a new topic of conversation). Converging results show that DMN spatial activity patterns for a given event are correlated across different individuals when they view a common movie stimulus^19–23^; a stable spatial pattern of brain activity (a neural state) is thought to persist across the duration of a perceived event^24–26^ and is similar across individuals. A few studies also suggest the presence of spatially organized brain responses during recollection. DMN patterns were correlated across people as they verbally recounted their memories of a recently-viewed movie, event-by-event^20,23^. Another study showed that activity pattern templates, generated by averaging movie-viewing data across subjects, could be used to identify reinstated event-specific patterns for each subject during spoken recall^27^. In these prior studies, posterior midline areas exhibited the strongest cross-subject reinstatement effects; at the same time, another study revealed that anterior midline DMN areas (e.g., mPFC) show relatively idiosyncratic activity during movie-viewing, indicated by low spatial correlations across individuals^28^. Thus, the existence of spatially organized DMN responses is supported by observations that activity patterns are correlated across different individuals when they view or remember a common event, with stronger cross-subject consistency observed in posterior than anterior midline aspects of the DMN. However, it remains unknown which features of episodic memories control this spatial organization of their concomitant neural responses.

For early sensory cortices, functional mapping typically relies on presenting tightly controlled stimuli which parametrically sweep feature space. For example, retinotopic maps in visual cortex are generated using expanding rings and rotating wedges, while tonotopic maps in auditory cortex are generated using ascending and descending tone patterns. How can we identify spatially organized responses in the posterior midline DMN which support the invariant representations underlying remembering, without knowing the relevant feature dimensions? One approach is to present a wide variety of complex stimuli, eliciting a wide variety of recollected content, and use data discovery methods to extract regularities in the resulting brain responses. Given the body of work showing that midline DMN neural representations are intimately linked to event and situation information^29^, it is crucial that both the encoding stimuli and the behavior at recollection encompass such content as well. With these data in hand, we can then search for neural activity patterns with certain properties that would be expected of spatially organized functional responses relevant to episodic recollection.

First, neural activity patterns should be stable across individuals, indicating a consistent physical organization in the brain; second, they should be linked between encoding and recall, indicating relevance for remembering; and third, they should occur repeatedly across the data recording session, as repetition enables the prediction of neural states from recurring features in the stimulus or behavior, while non-repeating states cannot be modeled in this way and are more difficult to distinguish from noise. Interestingly, inspection of group-averaged posterior medial DMN data during naturalistic movie-viewing suggests that indeed, subject-shared activity patterns appear to repeat across events; they also seem to fluctuate at timescales shorter than an event (Fig. 1a-b). These observations are not mutually exclusive with the notion of treating events as largely stable activity patterns^24,25^, but they do suggest that spatially-organized responses in the posterior medial DMN are not limited to a single state per event during movie-viewing, nor during recollection of such material.

**Fig. 1.**
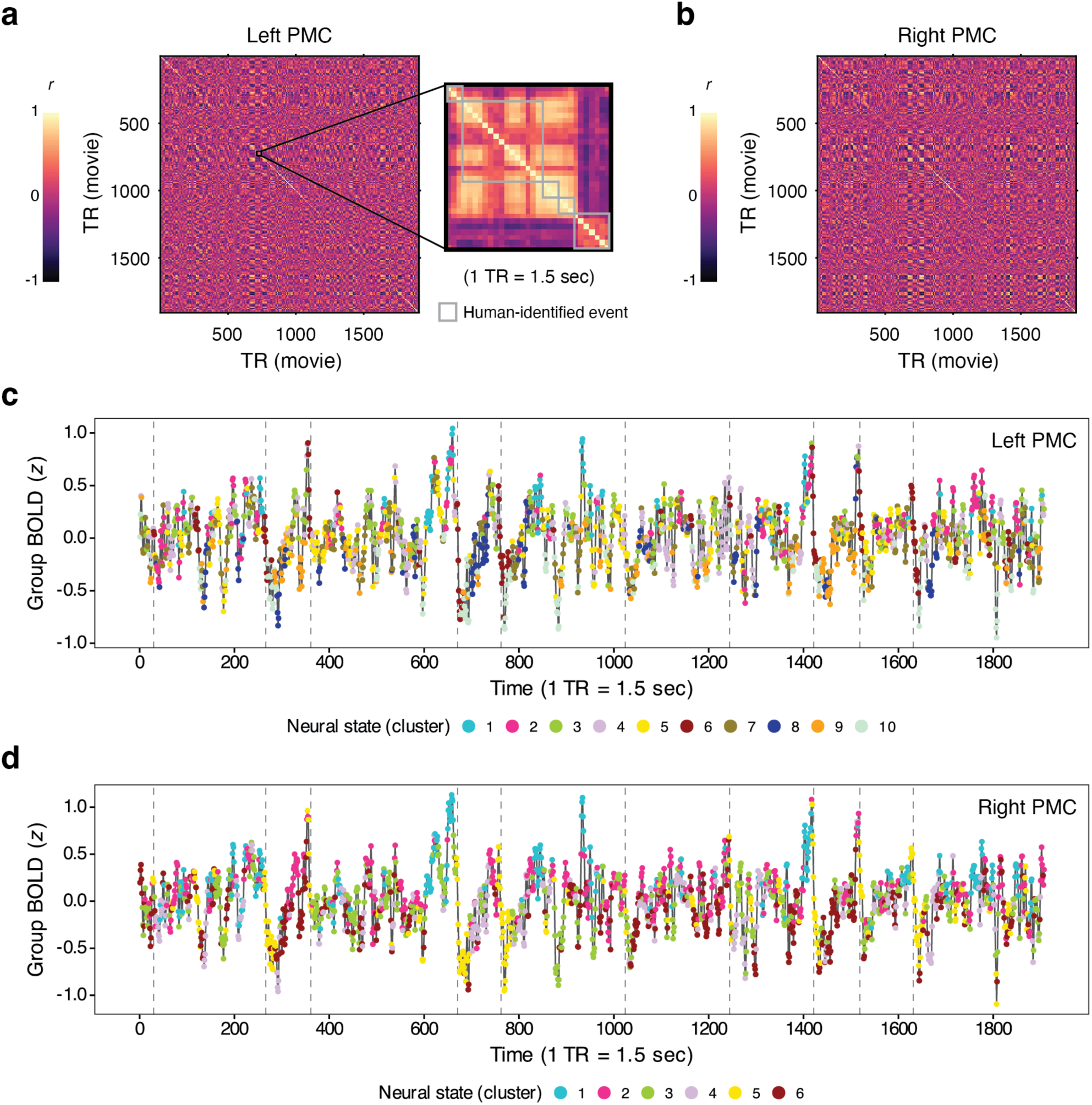
Group-averaged time-time correlation matrix during movie-viewing and neural state identification using *k*-means clustering. **a** We plotted a similarity matrix in which the correlation between pairs of multi-voxel patterns is illustrated over time, using group-averaged brain data from the left PMC during movie-viewing. Visual inspection indicates that subject-shared activity patterns associated with human-identified events are neither unique or entirely stable within an event. The yellow blobs in the off-diagonal sections of the matrix indicate that multi-voxel patterns are highly similar between pairs of time points (TRs) from distinct events as well as different movies. The data from all time points are shown in the matrix, totaling 1,907 TRs from two movie-viewing runs. **b** The similarity matrix for the right PMC is illustrated. **c** Individual time points from the group-average movie-viewing brain data were labeled by neural states. The BOLD time series in each graph was generated by averaging the activity across vertices in the group-average movie-viewing data. Neural states were identified based on the similarity of multi-voxel patterns at individual time points using *k*-means clustering. Each subject’s movie-viewing brain data was initially normalized (i.e., *z*-scored) across time within each functional run and then concatenated across the two movie-viewing runs. Colored dots indicate the cluster (i.e., neural state) membership of movie-viewing time points after applying *k*-means clustering. Gray dashed vertical lines indicate the onset of the first actual movie event for each movie, following its title scene. Results for *k* = 10 are shown for illustrative purposes for the left PMC data. **d** Neural states identified for the right PMC, with results for *k* = 6 shown for illustrative purposes.

In this study, we sought to elucidate the spatial organization of posterior midline DMN responses during episodic memory encoding and recall. We use a data-driven approach to discover a core set of neural states with three key features: 1) spatially aligned across individuals; 2) associated with reinstatement of naturalistic event information; and 3) repeated appearance across a wide variety of stimuli. We focus primarily on the posterior medial cortex (PMC), a key hub of the DMN which evinces robust reinstatement (i.e., reactivation) effects across numerous studies^20,23,30–35^, and which is thought to carry high-level information about complex events and situations^16,29^. Analyses were performed on an open neuroimaging dataset^36^ collected while subjects watched 10 short movies with widely varying plots, affective range, and visual styles, then recalled extensive details of the movies aloud from memory. In order to isolate signals with a common spatial organization across individuals, we averaged the movie-viewing data across brains; neural states were identified in these shared activity patterns via *k*-means clustering of time points. We then specifically selected neural states which re-appeared during recollection: we performed a modified reinstatement analysis in which movie-viewing time points were replaced with neural state patterns, and compared these to individual subjects’ recall data event-by-event. The analysis revealed that 16 neural states were sufficient to achieve memory reinstatement effects in PMC comparable to those reported using the established methods of the field. Consistent with prior studies, individual movie events tended to be dominated by a single neural state, though with substantial time also spent in other states. Stimulus-based models showed that the neural states were associated with actions and social-affective features in the movies. Overall, these findings enhance our understanding of the spatially organized functional responses within the posterior midline DMN which code for abstract event information during naturalistic encoding and recall.

## Results

We aimed to elucidate the structure of spatial response patterns underlying episodic recall of real-world events. Thus, we focused on the posterior medial portion of the DMN, which has been proposed to play a central role in representing situation models: internal mental models of entities, spatial context, actions and their relationships^16,37^. Representations of situation models are thought to be reactivated during recollection of real-world or narrative events, a notion supported by studies showing that activity patterns in PMC are significantly similar between encoding and recall of such events^20,22,27,30,33,34^. In other words, neural states during encoding are reinstated during remembering.

### DMN cortical areas, including posterior medial cortex, exhibited reactivation during spoken recall

Subjects (*N* = 15) viewed ten audiovisual movies, then described the movie events aloud from memory during fMRI scanning^36^. To confirm that posterior midline DMN areas exhibited reactivation while subjects verbally recalled the movie events, we first conducted a whole-brain reinstatement analysis following previously established protocols for naturalistic movie-viewing and recall data (e.g., ref.^19,20,23,27,38^). For both encoding (movie-viewing) and spoken recall data, events were defined according to human judgment (labels included with the fMRI dataset). Non-movie periods were excluded, leaving 190 events (see *Event segmentation* in Methods). Next, time series of multi-voxel patterns were obtained for a total of 400 cortical parcels drawn from an atlas derived from resting-state functional connectivity patterns^39^. While we use the term “multi-voxel” throughout this paper for consistency with common usage in the field, the analyses were conducted using vertices from the cortical surface. Within-subject reinstatement (i.e., encoding-retrieval similarity) analysis was performed wherein each subject’s brain activity pattern for each movie and recall event (i.e., the event pattern) was computed by averaging multi-voxel patterns across the time points corresponding to the event, then correlations were calculated between matching movie and recall event patterns. To assess the significance of the observed reinstatement effects, we constructed a null distribution by repeatedly shuffling the labels of movie events and calculating the similarity between the activity patterns from the scrambled movie events and intact recall events for 10,000 times. A two-sided *p*-value for the actual reinstatement was then calculated from the constructed null distribution for each cortical parcel. As in prior studies^20,23,27^, reactivation effects were observed throughout the DMN, and of particular interest for our study, in PMC (see Supplementary Fig. 1). For subsequent analyses, we defined the PMC ROIs as all parcels in the posterior medial portion of the DMN network, separately for the left and right hemispheres (see Supplementary Table 1 for ROI definition).

### Neural states were identified from movie-viewing data using a data-driven approach

Our next goal was to identify a set of neural states during movie-viewing that would be reactivated during later recall, regardless of their correspondence to individual movie events. While prior work has modeled brain activity states during movie-viewing as persisting across the duration of each event^24^, we wished to allow for neural states to be shorter than an event and to repeat across events. Thus, we used *k*-means clustering to group individual time points of the movie-viewing brain data according to the similarity of their spatial patterns. We chose not to apply dimensionality reduction, such as PCA or ICA, in order to retain the original multi-voxel pattern information and maintain comparability to the event-averaging methods used by prior studies^20,22,23^. The movie-viewing data were first averaged across all subjects to create a group-average vertex-by-time data matrix for PMC. Prior to clustering, the mean across the ROI was removed at every time point, so that the clusters would reflect the similarity of spatial patterns rather than the mean across vertices. To provide a range of clustering solutions which could later be compared to the recall data, *k*-means clustering was performed for *k*-values ranging from 2 to 20, separately for each hemisphere. Fig. 1c shows the outcome of *k*-means clustering applied to the left PMC data when *k*-value was 10 (see Fig. 1d for the right PMC data when *k* = 6). Note that the cluster numbers (i.e., labels) initially assigned by the *k*-means algorithm are random across iterations. Therefore, we reassigned cluster labels ordered by the magnitude of the average BOLD signal within each cluster after *k*-means clustering. For example, we labeled the cluster with the highest mean BOLD signal as cluster number 1 (Supplementary Fig. 2).

### How many neural states are needed to replicate standard reinstatement effects?

In order to determine how many neural states were needed to replicate standard reinstatement effects in PMC, we performed a modified version of a standard reinstatement analysis, wherein the original multi-voxel patterns from individual movie time points were replaced with neural state data from *k*-means clustering and then compared to recall data (Fig. 2a; see *Modified reinstatement analysis* in Methods). Specifically, for each value of *k* (2 to 20), a cluster pattern was created for every *k*-means-derived group of time points by averaging across all time points assigned to that cluster. For example, when *k* = 6, there were 6 different cluster patterns. Each time point in the movie data was replaced with the cluster pattern matching its cluster group. If time point 100 was assigned to cluster 5, time point 100 was replaced with cluster pattern 5 (the average of all time points identified as cluster 5). Importantly, no alterations were made to the recall data. The brain activity pattern for each movie and recall event was computed by averaging multi-voxel patterns across the time points corresponding to the event, and then correlations were calculated between matching movie and recall event patterns. Note that movie event patterns were calculated using cluster patterns obtained from group-average data (Fig. 2a, right), whereas recall event patterns were computed from each subject’s own spoken recall data. In other words, the calculation of recall event patterns remained identical between the standard and the modified version of the reinstatement analyses. The modified reinstatement analysis was repeated 10 times, with *k*-means clustering re-run in each iteration to assess the reliability of the results. As shown by the gray lines in Fig. 2b-c, the clustering outcomes and the computed reinstatement effects were highly similar across iterations, indicating consistency in the overall pattern.

**Fig. 2.**
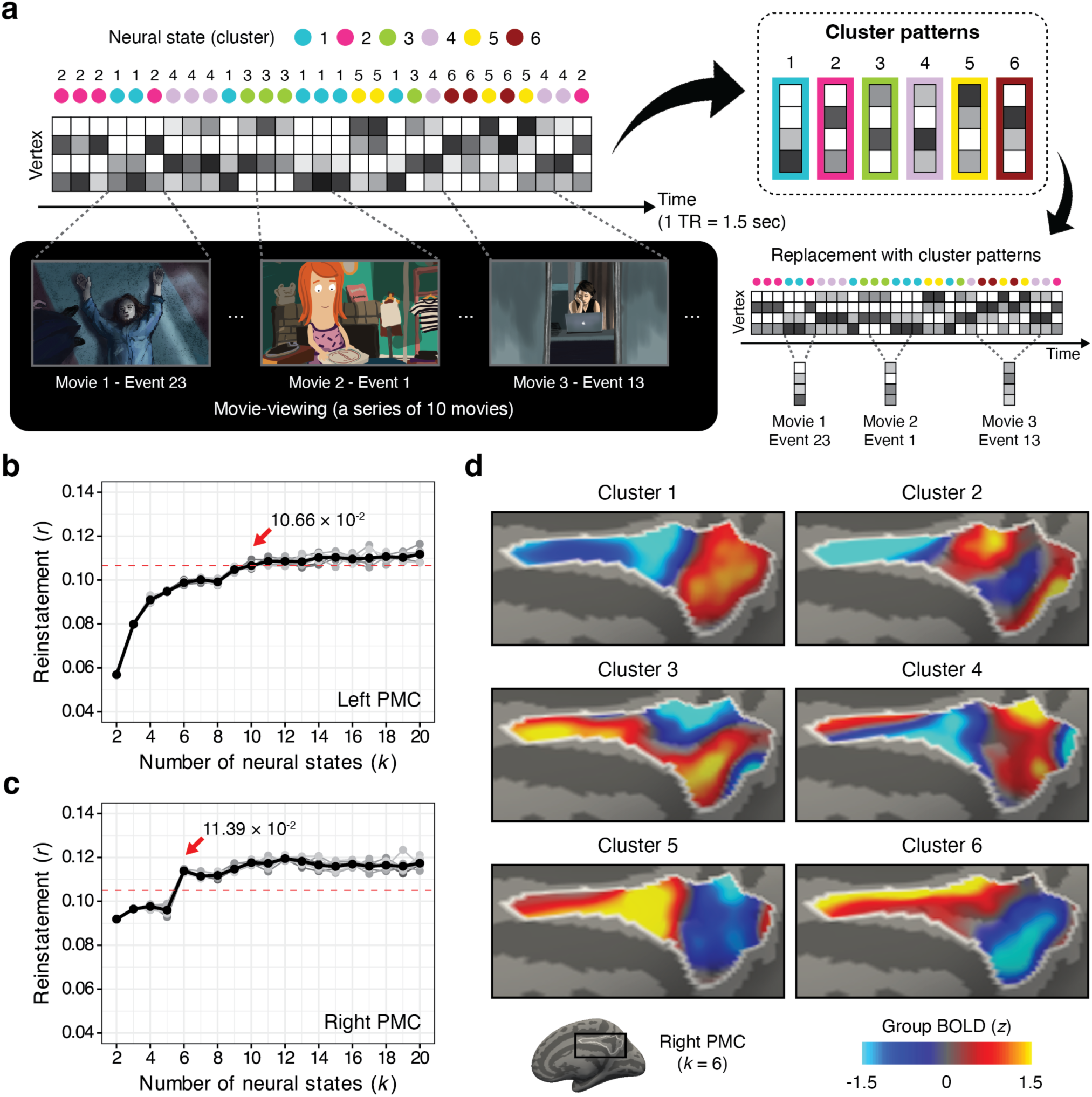
Memory reinstatement effects achieved by PMC component neural states. **a** A modified version of a reinstatement analysis was performed. The analysis procedures are illustrated for *k* = 6. Six neural states were identified based on the similarity of spatial activity patterns across individual time points in the group-average movie data (see Fig. 1c and Fig. 1d for the actual *k*-means clustering outcome for the left and right PMC). A cluster pattern, reflecting the typical activation pattern of each neural state, was created by averaging the activation patterns across time points labeled with the same neural state. Movie-viewing data were re-generated by replacing the activity patterns at each time point in the original movie brain data with the corresponding cluster pattern. Movie event patterns, defined as the spatial activity patterns averaged across the duration of a movie event, were then calculated using the newly generated movie-viewing data. **b-c** In each plot, the red dashed line illustrates 95% of the standard reinstatement effects observed in each region: 10.65 × 10^−2^ for the left PMC and 10.50 × 10^−2^ for the right PMC. Based on results averaged across iterations (indicated by the black bold line), we selected the minimum number of neural states required to achieve more than 95% of the standard reinstatement effects in the dataset for each region. As a result, 10 neural states were chosen for the left PMC and 6 neural states for the right PMC. The red arrow indicates the memory reinstatement strength value from the iteration-averaged curve for the selected number of neural states in each hemisphere (achieved reinstatement strength: 10.66 × 10^−2^ for the left PMC when *k* = 10 and 11.39 × 10^−2^ for the right PMC when *k* = 6). Gray shades show the results from each iteration (out of 10) of *k*-means clustering. **d** The cluster patterns of 6 neural states identified in the right PMC are visualized on the cortical surface. The results from one iteration of *k*-means clustering are presented for visualization purposes (the same data shown in Fig. 1d). For visualization, the vertex values of each cluster pattern were *z*-scored within the ROI.

**Fig. 3.**
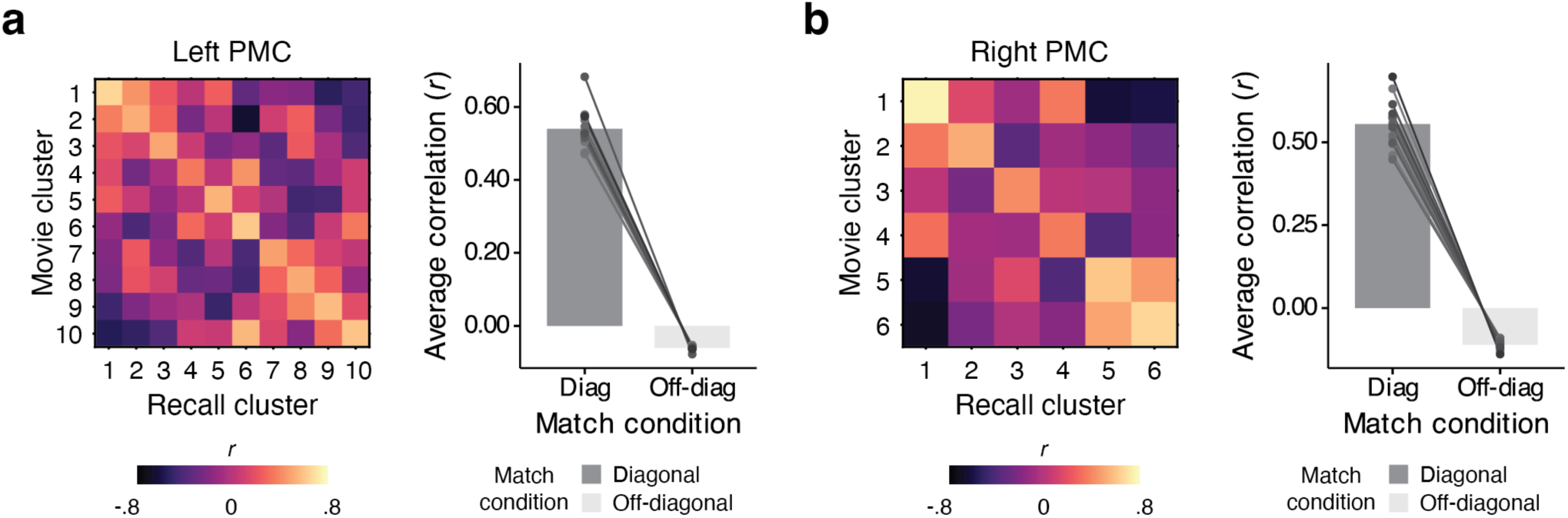
Correspondence of movie cluster patterns with recall cluster patterns in the left and right PMC. **a** In addition to the 10 “movie” cluster patterns identified for the left PMC, we performed *k*-means clustering on each subject’s free spoken recall data to obtain 10 “recall” cluster patterns per individuals. For each subject, we first identified the best correspondence matrix by reordering the labels of an individual’s recall cluster patterns to maximize the average diagonal of the movie-recall cluster-cluster correlation matrix in all possible cases. The matrix shown here illustrates the group-average of individuals’ best match matrices. The line graphs show the average correlation in the diagonal and off-diagonal elements of each subject’s best correspondence matrix, indicating a consistent pattern across subjects. The bar graphs show the values averaged across subjects. **b** The same analysis was performed for the right PMC, where six recall cluster patterns were identified. Diag: Diagonal, Off-diag: Off-diagonal.

We found that, in the left PMC, 10 neural states were sufficient to achieve 95% of the reinstatement value from the original (standard) analysis, based on results averaged across iterations (Fig. 2b). In the right PMC, 6 neural states were sufficient (Fig. 2c; see Supplementary Fig. 3 for the results of the early auditory and early visual cortical areas for comparison). Individual events were composed of multiple neural states, and individual neural states were found in multiple movies. Thus, despite the inherent variability in individual’s recall utterances and descriptions, memory reactivation was explained by a relatively small number of neural states identified at the group level. Fig. 2d shows the cluster patterns of 6 neural states identified in the right PMC, visualized on the brain surface. The results reported in the following sections make use of only one (of the 10) *k*-means clustering solutions. Supplementary Fig. 4 illustrates the cluster patterns identified in the left PMC when *k* = 10.

See Supplementary Fig. 5 for similar analyses in the angular gyrus. Even with 20 states in the left angular gyrus, 95% of the standard reinstatement effects were not reliably achieved. As in PMC, fewer neural states (11 states) were sufficient for the right angular gyrus.

### Neural states identified during movie-viewing are also present during verbal recollection

To determine whether the neural states identified during movie-viewing reappear during recall, we applied *k*-means clustering to each subject’s recall data, grouping time points to generate recall cluster patterns. We then identified labels for each subject’s recall cluster patterns that best matched the previously identified movie cluster patterns, based on maximizing the average diagonal of the movie-by-recall cluster correlation matrix (see *Identification of neural states: Recall* in Methods). This was achieved by reordering each subject’s recall cluster patterns while preserving the original order of movie cluster patterns. After applying the best match procedures, we obtained the group-average correlation matrix by averaging across individuals. We evaluated the significance of this match using a permutation test, wherein the order of recall cluster patterns was randomly shuffled for each subject before calculating a new group-average correlation matrix.

Fig. 3 illustrates the group-average correlation matrix of the best correspondence of movie and recall cluster patterns (average diagonal = .54, average off-diagonal = -.06 in the left PMC; average diagonal = .56, average off-diagonal = -.11 in the right PMC). The analysis was performed for *k* = 10 (left PMC) and *k* = 6 (right PMC), as determined in the prior analysis (see Results: *How many neural states are needed to replicate standard reinstatement effects?)*. The observed average diagonal in the group correlation matrix was significantly different from the null distribution (two-sided test; mean of the null distribution of the average diagonal = −1.27 × 10^−4^, *p* < .0001 in the left PMC; mean of the null distribution of the average diagonal = −3.89 × 10^−4^, *p* < .0001 in the right PMC; Supplementary Fig. 6). These results suggest that a similar set of cluster patterns (i.e., neural states) is shared between the stimulus-driven (i.e., movie-viewing) and internally driven (i.e., verbal recall) processes.

### Separation, distribution, and duration of the neural states

Previously, we observed that neural states identified through *k*-means clustering repeatedly occurred across movies, as confirmed by visual inspection (Fig. 1). Interestingly, the duration of neural states tended to be shorter than the human-defined movie events and were distributed across different movies, as well as across different events within a single movie. Next, we aimed to characterize the properties of the PMC neural states we identified. As an overview, we observed that 1) similar spatial patterns of brain activity were evoked by movie scenes with different content, and 2) despite the presence of transient neural states, individual movie events tended to be dominated by a single PMC neural state, with different events dominated by different states.

#### Cluster separation

To what extent were the PMC individual time point multi-voxel patterns similar within a cluster and distinct between clusters? Previously, we constructed a time-by-time correlation matrix of multi-voxel patterns for every pair of time points in the group-average movie-viewing data (Fig. 1a-b). This similarity matrix was sorted based on clusters rather than chronological order (Fig. 4a-b). We then calculated the average correlations for TRs within and across clusters. As expected given the *k*-means clustering procedure, individual time point multi-voxel patterns were positively correlated within clusters and less correlated between clusters (within-cluster similarity averaged across clusters = .38, between-cluster similarity averaged across cluster pairs = -.04 in the left PMC; within-cluster similarity averaged across clusters = .33, between-cluster similarity averaged across cluster pairs = -.05 in the right PMC). Self-correlations (i.e., multi-voxel pattern correlations of the same TR, which should equal 1) were excluded from the calculation of average within-cluster similarity by considering only the upper triangle of each sorted matrix. Visual inspection indicates that the clusters were not completely uncorrelated, which is expected since orthogonality is not a requirement of *k*-means clustering.

**Fig. 4.**
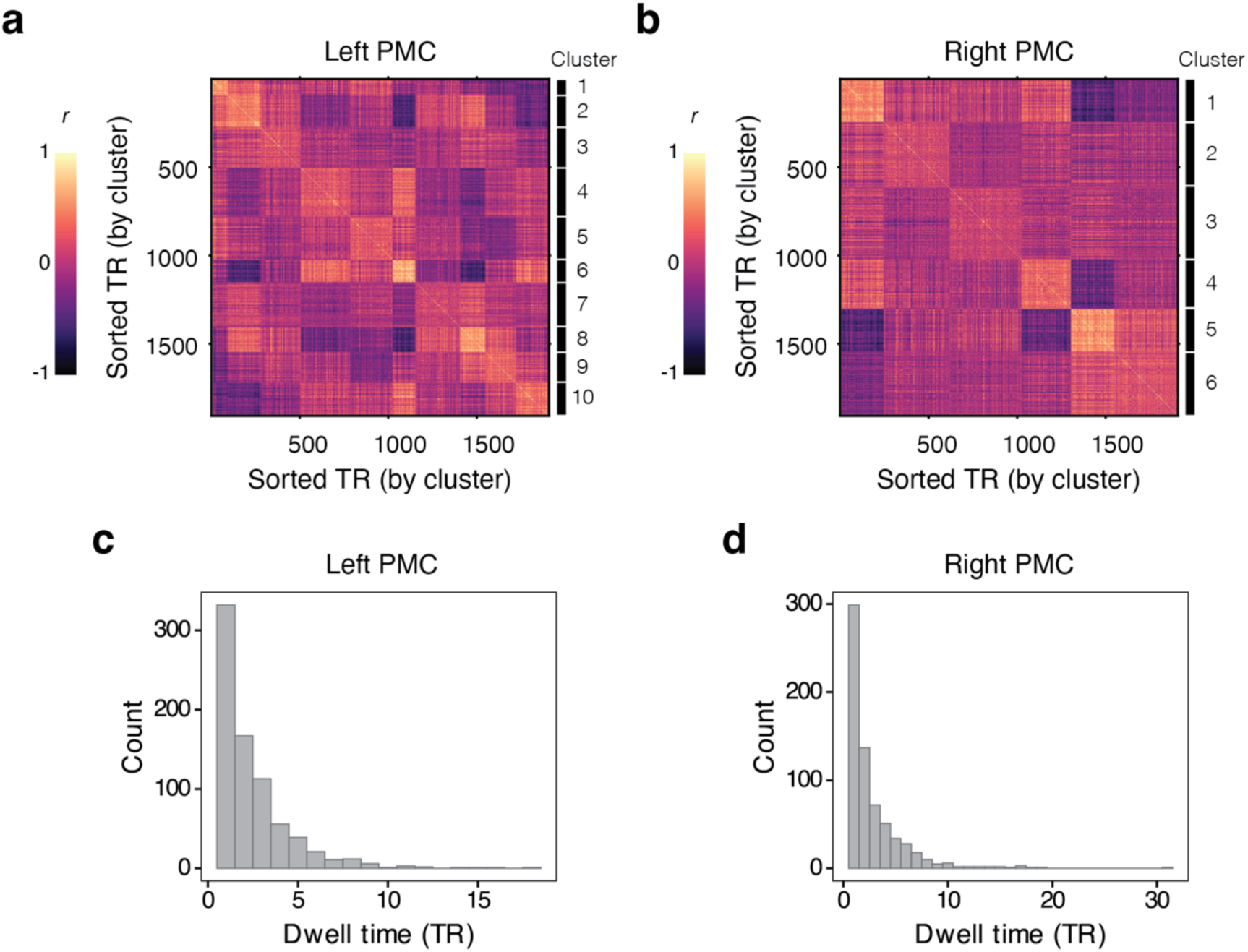
Cluster separation and duration of neural states during movie viewing. The time-time correlation matrix for movie-viewing data was sorted by clusters. TRs grouped into the same cluster (i.e., neural state) are shown as yellow blobs, indicating high similarity between pairs of TRs; 1 TR = 1.5 sec. **a** In the left PMC, *k* = 10 was selected in the preceding modified version of the reinstatement analysis. The first yellow blob in the upper-left corner corresponds to cluster 1, while the blob in the bottom-right corner shows cluster 10. **b** Similarly, in the right PMC, clusters appear along the diagonal from the top left (cluster 1) to the bottom right (cluster 6) for the selected *k*-value (*k* = 6). **c-d** Histograms of neural state dwell time are shown for **c** the left PMC and **d** the right PMC.

#### Distribution of neural states across movies

Did the time points assigned to a given cluster predominantly correspond to a single movie, or were they distributed across movies? We calculated the number of TRs in each cluster that appeared across ten movies (i.e., their occurrences). In the left PMC, 8 of the 10 neural states occurred at least once in all ten movies. In the right PMC, all 6 neural states occurred at least once in all ten movies. Supplementary Figs. 7-8 illustrate the occurrences of each neural state in each movie.

#### Duration of neural states

In prior studies, individual events were modeled as having a stable, persistent activity pattern across the duration of the event^24,25^. To assess whether this holds for the current PMC neural states, we examined their temporal stability. We defined a “cluster segment” as any consecutive series of time points assigned to the same cluster in the group-average movie-viewing data; “dwell time” was defined as the duration of individual cluster segments. Fig. 4c illustrates the distribution of dwell times in the left PMC, which ranged from 1 to 18 TRs (1.5 – 27 sec) and peaked at 1 TR. In the right PMC, dwell time ranged from 1 TR to 31 TRs (1.5 – 46.5 sec), also peaking at 1 TR (Fig. 4d). We observed a cluster segment in the right PMC with a dwell time of 31 TRs at the beginning of Movie 5 (“Keith Reynolds”), making it the only cluster segment in the dataset that exceeded the longest movie event duration (41 sec). The distributions of dwell times across neural states are illustrated in Supplementary Figs. 9-10. Overall, neural state dwell times were mostly transient and shorter than human-labeled events, but occasionally were as long as movie events (mean movie event duration across the 10 movies ranged from 9.4 to 18.4 sec).

#### Individual movie events were mostly dominated by a single neural state

Based on prior studies that revealed stable multi-voxel patterns during the duration of movie events in the DMN^24,25^, we predicted that individual movie events would predominantly consist of a single neural state, despite the transient nature of neural states and their distribution across different movies. We calculated the percentage of time points assigned to each cluster for each movie event and identified the *dominant* neural state (the one with the largest percentage, regardless of contiguity). For both the left and right PMC, we found that the dominant neural state was present for more than half the event duration, on average, at the *k*-values selected by the preceding modified reinstatement analysis (56.46% for the left PMC when *k* = 10; 65.31% for the right PMC when *k* = 6). This observation is compatible with prior characterizations that stable PMC states are associated with individual movie events, though it suggests that each event is also composed of multiple transient neural states.

#### Reinstatement analysis using dominant clusters alone

Having observed that a dominant neural state composes individual movie events, we conducted an alternative version of the modified reinstatement analysis, primarily using the dominant neural states for movie event patterns. For each movie event, the event pattern was replaced with its corresponding dominant neural state (i.e., cluster pattern), while each subject’s own recall event patterns remained unchanged. The analysis was performed using the previously selected *k*-values (i.e., 10 cluster patterns for the left PMC and 6 cluster patterns for the right PMC), based on a single iteration of clustering; the same clustering outcome reported in other figures. The observed reinstatement strength fell below 95% of the standard reinstatement effects (reinstatement strength: 8.91 × 10^−2^, 79.5% of its standard effects in the left PMC; reinstatement strength: 9.12 × 10^−2^, 82.5% of its standard effects in the right PMC), contrasting with the results in Fig. 2b-c, in which the full set of cluster patterns was used. These findings indicate that a dominant cluster alone was not sufficient to achieve the standard reinstatement effects.

### Neural states were associated with actions and social-affective features in the movies

Memory reactivation in cortical areas has been suggested to reflect the retrieval of information from past experiences^20,24,40–43^. What type of information is associated with the PMC neural states we identified? To address this question, we performed a voxel-wise encoding model analysis^44,45^ to test whether a set of neural states could be predicted from movie features. We constructed three models from movie features: 1) an “action” model, as research has shown that perceiving physical changes in actions is critical for event boundary judgments in videos of everyday activities^46,47^, and the retrieval of autobiographical memories is facilitated by knowledge of stereotyped sequences of actions in events^48^; 2) a “social-affective” model, as social-affective features of actions in naturalistic scenes have been shown to be processed at relatively late stages of cortical temporal processing^49^. Additionally, DMN areas with long information processing timescales^50^, such as PMC, are hypothesized to receive input from lower-level sensory processing areas during natural narrative processing^51^; and 3) a full model, which includes both actions and social-affective features as predictors. Action labels were obtained from 309 online participants who watched the movies and generated text descriptions of the actions, then converted to word embedding vectors for analysis (Fig. 5a). The same participants rated social-affective properties (sociality, valence and arousal) at each time point of the movie (see *Actions and social-affective features* in Methods). Model prediction performance was evaluated using a leave-one-TR-out cross-validation approach (Fig. 5b). We examined whether the spatial activation pattern predicted by the model at a given TR was most similar to the cluster pattern corresponding to the held-out TR. Cluster patterns were newly created in each iteration, excluding the held-out TR (see *Voxel-wise encoding model estimation and validation* in Methods).

**Fig. 5.**
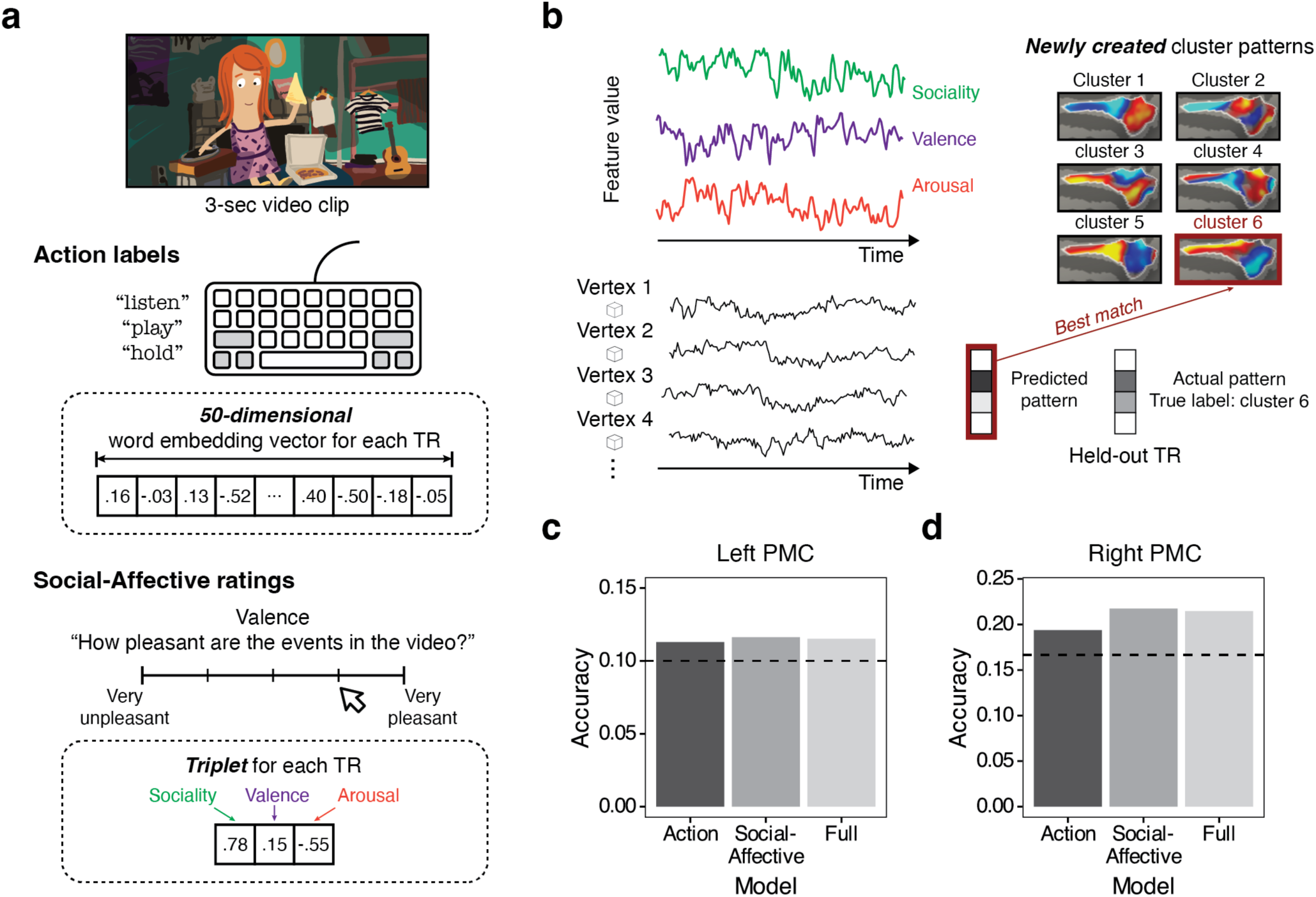
Encoding model analysis results. All three models showed above-chance predictions for both hemispheres. **a** Online participants provided action verbs and rated the social-affective properties of each 3-second video clip. For each TR (1.5 sec), a 50-dimensional GloVe^52^ word embedding vector was generated by averaging the vectors across verbs. Additionally, a triplet of subject-average ratings for sociality, valence, and arousal was calculated for each TR. These movie features were used in an encoding model. **b** Each model’s predictions were evaluated using a leave-one-TR out cross-validation scheme. We tested whether the predicted pattern of a held-out TR corresponded best to the cluster of its actual pattern. Cluster patterns were recalculated for each of the 1,780 iterations (TRs). **c** In the left PMC, the chance level was .1 (1/10), as there were 10 neural states that each model had to predict from movie features for the chosen *k* = 10. **d** In the right PMC, the chance level was .167 (1/6), when *k* = 6. The gray-shaded bars correspond to the different encoding models we constructed.

We found that all three models predicted PMC cluster patterns above chance for the chosen number of neural states in the preceding modified reinstatement analysis (action model accuracy = .113, social-affective model accuracy = .116, full model accuracy = .115 in the left PMC when *k* = 10; action model accuracy = .194, social-affective model accuracy = .217, full model accuracy = .215 in the right PMC when *k* = 6; Fig. 5c-d). The social-affective model showed numerically higher accuracy than the action model. Additionally, the accuracy of the full model was numerically lower than that of the social-affective model. This is likely due to the high correlation between actions and social-affective features, as the action verbs generated by online participants often included actions socially relevant actions (e.g., greet and converse) rather than being confined to physical motions (e.g., walk and throw). It should also be noted that three models also predicted cluster patterns in early auditory and visual cortices (Supplementary Figs. 11-12). Thus, while actions and social-affective features predicted neural state activity patterns in PMC, this relationship was not limited to high-order cortical areas.

### Some neural states were strongly correlated with a previously reported generalized boundary pattern

Next, we investigated whether any of the PMC neural states identified in our analyses corresponded to a previously reported “generalized boundary pattern” seen at major context boundaries (i.e., between movies) during both encoding and recall^53^. We examined the frequency of each PMC neural state during movie transition periods, which were defined as the 15 seconds (10 TRs) immediately following the end of the last event in each movie. As illustrated in Fig. 6, we found a specific neural state that appeared predominantly during movie offset periods (cluster 6 in the left PMC and cluster 5 in the right PMC).

**Fig. 6.**
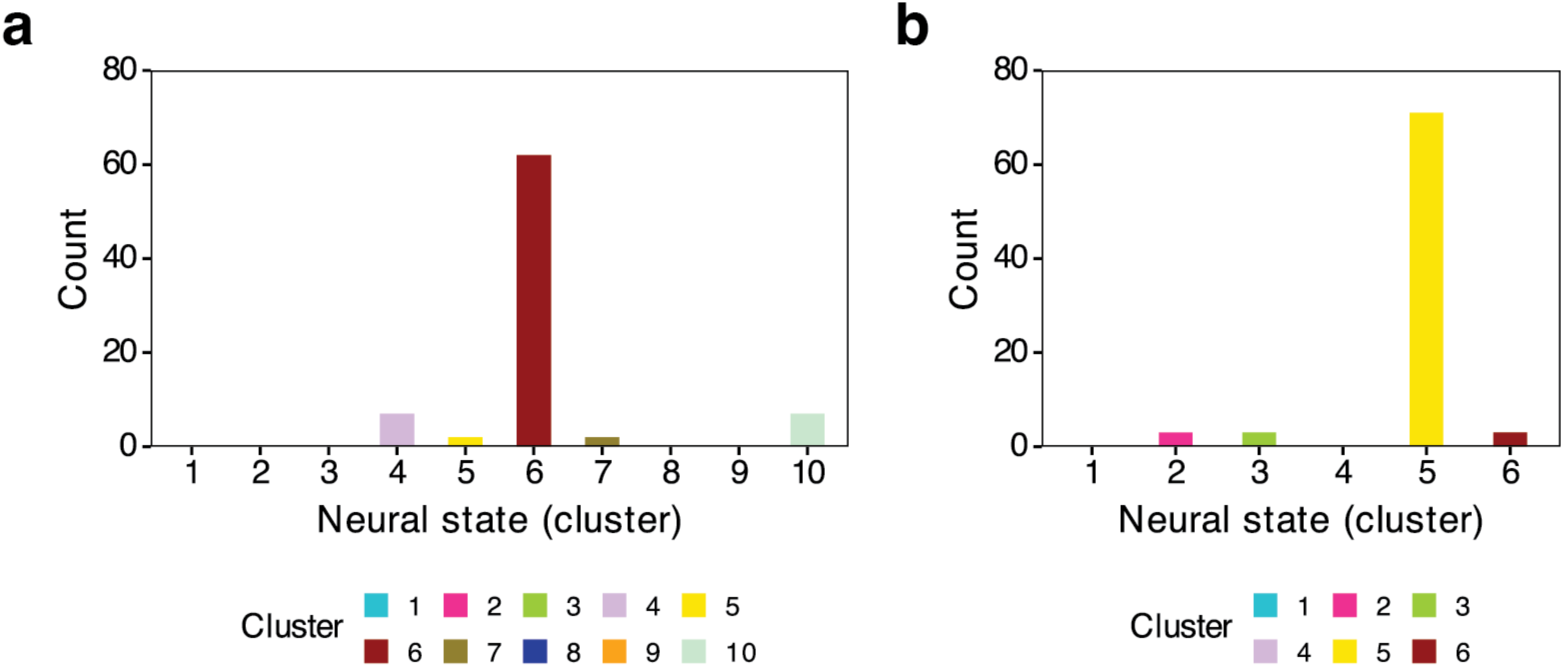
Neural states that most frequently appeared during movie transitions in the left and right PMC. The figure illustrates the occurrences of each neural state during movie transitions. Due to the limited number of available time points, the last movie in each movie-viewing run was excluded, resulting in a total of 80 TRs from eight movies. The cluster labels are based on one iteration of the *k*-means clustering outcome used for other figures in this paper. Across iterations, one neural state consistently appeared most frequently during movie transition periods. For example, in the left PMC, cluster 6 appeared most frequently in three out of 10 iterations. It is important to note that the initial cluster labels assigned by the algorithm do not follow any inherent organizational principle. Instead, the labels were organized based on the within-cluster averaged BOLD signals. **a** Left PMC. **b** Right PMC.

We further examined whether the cluster patterns were most strongly correlated with a spatial activation pattern at movie boundaries during recall. We calculated a group-average recall boundary pattern by averaging recall boundary patterns across subjects. For each subject, a recall boundary pattern was calculated by first averaging the multi-voxel patterns during the 15 seconds (10 TRs) following the completion of the recall of a previous movie and then averaging these patterns across all movie recall transitions. As expected, cluster pattern 6 (from the movie data) was most strongly correlated with the group-average recall boundary pattern in the left PMC (*r* = .91), whereas the strongest correlation between cluster pattern 5 (from the movie data) and the group-average recall boundary pattern was found in the right PMC (*r* = .95); see Supplementary Fig. 4 for a visualization of cluster pattern 6 in the left PMC and Fig. 2d for cluster pattern 5 in the right PMC. We additionally performed an analysis in which cluster patterns were calculated using each individual subject’s movie-viewing data, applying the clustering solution and cluster labels from the group data. Consistent with the preceding analysis, in which group-average movie-data were used, the strongest correlation was found for cluster pattern 6 in the left PMC (*r* = .72) and for cluster pattern 5 in the right PMC (*r* = .78), averaged across subjects (Supplementary Fig. 13). Together, these results show that two PMC neural states, one in each hemisphere, were strongly associated with movie transitions during both encoding and recall. These may reflect a cognitive state of mental context flushing, as proposed by prior research^53^.

## Discussion

In this study, we identified a core set of neural states in PMC that underlie reinstatement of naturalistic event memories during spoken recollection. PMC has been suggested as a core hub within the posterior midline DMN, consistently showing strong memory reinstatement effects in numerous prior studies using both constrained and naturalistic stimuli^20,23,30–35^. The PMC neural states were first derived from group-level movie-viewing data, capturing stimulus-locked brain signals reliably evoked by 10 different movies across subjects. We found that only a small number of states was needed in order to achieve the same level of movie-recall reinstatement effects in PMC (*k* = 10 in the left PMC and *k* = 6 in the right PMC) that are commonly reported in the literature. We further demonstrated that the neural states extracted during movie-viewing could also be identified in each individual subject’s recall data. These results provide evidence for the existence of a spatially organized neural code in PMC that is shared across individuals and evoked during recall of complex, naturalistic experiences. Additionally, we characterized several properties of these neural states. While individual movie events were associated with multiple neural states, one state tended to dominate each event, with the dominant state varying across events. Moreover, these neural states were predicted by actions and social-affective features in cinematic movies, as determined by human raters. Finally, one of the identified neural states corresponded well to a previously reported pattern observed at moments of major transitions between movies and during recall of movies^53^. In summary, these results support the idea that representations of remembered naturalistic events in PMC may be composed of a surprisingly small set of spatially consistent neural states.

To our knowledge, this is the first study to characterize the properties of spatially organized neural states underlying naturalistic recall. However, several prior studies have investigated whether brain signals evoked during naturalistic movie viewing can be characterized in a low-dimensional space. At the whole-brain level, large-scale neural dynamics can be characterized within a low-dimensional latent state space, derived from patterns of activity and covariance across brain areas. These latent states are associated with attentional engagement during movie-viewing^54^. Another study identified neural states simultaneously from resting-state and movie-viewing data, based on whole-brain patterns of activation across brain networks. These neural states covaried with physiological changes, such as heart rates and pupil dilation, and were linked to certain movie features^55^. Furthermore, large-scale functional organization of the DMN includes specialization of subregions; for example, multi-voxel patterns in the posterior midline DMN are similar for events encoded in the same spatial context, while patterns in the anterior DMN are similar for events involving the same agent^34^. Focusing in further on the posterior medial aspect of the DMN, one study found that brain activation in the medial parietal cortex is topographically organized by information types in memory, such as how familiar subjects thought the scenes in the videos were when watching a video of their own memories^56^. Another study, the only one to our knowledge that incorporates data from recall in the absence of stimuli, estimated that fewer than 20 dimensions of information were encoded in PMC event patterns shared across subjects during movie-viewing and recall^20^, though these were not constrained to be spatially conserved across individuals. More work is needed to more fully understand the dimensionality and topography of DMN signals during recollection.

Our results raise the possibility that distinct subnetworks within the DMN cooperatively contribute to the spatial and functional organization of PMC signals involved in naturalistic event memory encoding and retrieval. Prior work has revealed functionally distinct and spatially separate subnetworks within the canonical DMN by parcellating brain areas based on intrinsic functional connectivity. For example, studies have identified two interdigitated subnetworks, network A and network B, within the DMN at the individual level^57^. Network A, connected to the parahippocampal cortex, is primarily involved in episodic projections, whereas network B, linked to the temporoparietal junction, is strongly engaged in theory-of-mind tasks^58^. The DMN has also been segmented into three interconnected subnetworks focused on cortico-hippocampal interactions: the posterior medial (PM), anterior temporal (AT), and medial prefrontal (MP) networks^59^. In an earlier framework^16^, the PM network was proposed to play a crucial role in episodic tasks, such as remembering or imagining episodes (i.e., constructing mental models about “what is happening”; situation models). Recent work has further suggested heterogeneity within the PM network^60,61^. In this study, we focused on the posterior midline DMN, also referred to as the PM network. Specifically, we explored spatially organized brain activity patterns that serve as the “basis” for event memory reinstatement in PMC. One interesting observation from our study was that some PMC neural states showed a topographically opposite pattern of activation (e.g., cluster 1 and cluster 6 in the right PMC) as illustrated in Fig. 2d. Similar results were found for the neural states identified in the left PMC, as shown in Supplementary Fig. 4. In our study, the defined PMC ROI encompassed anatomical locations likely corresponding to the two distinct subnetworks, network A and network B, within the DMN. An intriguing possibility is that the topographic differences observed between these neural states reflect differential contributions of these two subnetworks during the encoding of naturalistic events. Future work could explore whether the PMC neural states we identified are associated with processing biased toward either episodic details or the inference of movie characters’ intentions, as well as the relative contributions of the two subnetworks to these biases.

While the overall duration of neural states (i.e., dwell time) was generally short, we found that movie events were predominantly associated with a single PMC neural state, though the dominant state varied across events. In other words, the presence of rapidly changing PMC neural states does not necessarily contradict the established claim in the field that multi-voxel patterns are stable within an event^24^. Prior studies have consistently demonstrated that spatial patterns of activation in DMN areas are relatively stable over time within an event and transiently change at moments corresponding to event boundaries. This has been shown in numerous studies through the application of latent state-detecting computational models to movie-viewing data in a data-driven manner^24,25,27,62,63^, as well as by directly measuring the magnitude of voxel pattern shifts according to event boundary strength using a sliding window approach anchored to human-identified event boundaries^64^. Based on our observations, PMC appears to represent movie events by integrating different sets of neural states, although the underlying rules remain unknown. Future studies could investigate the specific computations within PMC that generate event patterns with this small set of neural states.

The core PMC neural states investigated here could be intriguing in contexts beyond event memory. In addition to serving as building blocks for episodic representations in these high-level cortical areas, one interesting possibility is that they may be utilized in verbal communication between individuals. A clue comes from prior empirical research, which found that multi-voxel patterns in the DMN are similar across three groups of subjects: 1) a group watching a cinematic movie, 2) another group listening to audio of someone verbally recalling the movie, and 3) one subject recalling the movie themselves^23^. These results suggest that DMN neural codes not only support construction of episodic representations from memory, they can also be transmitted to other individuals via language. A recent review paper proposed that the DMN can be understood as a sense-making brain network that integrates extrinsic (i.e., sensory input driven) and intrinsic (e.g., long-term memories or beliefs) information^65^. Given that subject-shared responses in the DMN are associated with behaviorally measured shared interpretations and shared meanings of experimental stimuli^13,66–68^, the PMC neural states reported in this paper may support not only event memory but also verbal communication between individuals (see ref.^69–71^ discussing the intimate link between memory and communication).

In summary, we demonstrated that brain responses in high-level DMN areas can be effectively modeled as relatively low-dimensional, spatially organized activity patterns using a data-driven approach. Importantly, the spatial organization (i.e., neural states) we uncovered was shared across individuals, was sufficient for standard event memory reinstatement analyses, and was present during both naturalistic stimulus presentation (movie-viewing) and an internally oriented memory task in the absence of a stimulus. Additionally, we observed an association between the neural states and the information carried by the stimulus, specifically actions and social-affective features in movies. These findings highlight that spatially organized brain signals underlying event memory encoding and retrieval can be uncovered in the posterior midline DMN, despite the inherent complexity of real-world, naturalistic experiences.

## Methods

### fMRI data source

We analyzed a publicly available dataset^36^ in which 15 subjects (male = 5) watched 10 short films, then verbally recounted their memories of the movies, all during fMRI scanning. Informed written consent was provided by all subjects in accordance with procedures approved by the Princeton University Institutional Review Board.

### Stimuli

The experimental stimuli consisted of 10 short audiovisual movies (on average 4.54 minutes long, range 2.15 – 7.75 minutes). The movies varied widely in their content, narrative structures, emotion, visual styles and formats. For example, three of them were cartoons, whereas others were live-action movies. Each movie, prepended with a 3-sec to 6-sec title screen, was presented only once; the movies were presented in two scanning runs (five movies per run, 24.9 minutes and 22.9 minutes). A 39-sec introductory cartoon (“Let’s All Go to the Lobby”) was played at the beginning of each movie-viewing run.

See the prior study^36^ for detailed information about stimuli and experimental procedures.

### Event segmentation

We used timestamps provided by a prior study^36^. Movie title scenes and scenes from “Let’s All Go to the Lobby” were excluded, resulting in 202 events. For memory reinstatement analyses only, we further excluded 12 events which were recalled by fewer than five subjects, leaving a total of 190 events.

### Cortical parcellation and region of interest (ROI) definition

We performed a whole-brain analysis using an atlas consisting 400 cortical parcels (200 parcels from each hemisphere) derived from the resting-state functional connectivity (17 networks)^39^. Following the ROI definitions from prior studies (see ref.^22^ for the PMC and early visual cortex ROIs and ref.^72^ for the early auditory cortex ROI), we performed the ROI analysis on the PMC and early sensory processing areas, specifically the early visual and auditory cortices, as control regions. Supplementary Table 1 lists the parcels used for the ROIs.

### Whole-brain within-subject standard reinstatement analysis

A whole-brain reinstatement analysis was performed using each subject’s parcellated data, following established procedures for movie and recall data (e.g., ref.^20^). Movie and recall event patterns were calculated for each parcel by averaging multi-voxel patterns across time for each event. To measure within-subject reinstatement strength, we computed a correlation between each matching pair of movie and recall event patterns, then averaged across event pairs and subjects, resulting in one group-average correlation value per parcel. To test which cortical areas were significantly involved in memory reactivation, a permutation test was performed. A null distribution was constructed by shuffling movie event labels within each subject and recalculating the group-average correlation value 10,000 times^73^. The group-average value was compared against the null distribution and FDR-corrected across parcels (*q* = .05, two-tailed).

### Identification of neural states: Movie-viewing

Neural states in PMC were identified from group movie-viewing data using *k*-means clustering. We applied *k*-means clustering to all available time points, including periods corresponding to the movie title scenes and “Let’s All Go to the Lobby,” to identify distinct neural states. The *k*-means clustering algorithm partitioned data in an unsupervised manner into *k* clusters (i.e., *k* distinct neural states) based on the similarity of individual time-point multi-voxel patterns, measured as 1 - Euclidean distance. For example, if *k* = 10, the algorithm assigned each time point a cluster membership value between 1 and 10 (inclusive). We first obtained a time series of multi-voxel patterns averaged across all subjects, in order to identify shared signals across individuals. The mean BOLD signal, averaged across vertices, was subtracted from every vertex at each time point to minimize univariate effects on clustering. Clustering was performed for *k*-values ranging from 2 to 20, with 100 random initializations of cluster centers for each *k*. For each *k*-value, a set of “cluster patterns” was then generated by averaging the spatial activation patterns across time points grouped into the same cluster, regardless of their event or movie correspondence. Note that the cluster numbers (labels) initially assigned by the *k*-means clustering algorithm lack any inherent organizational principle. We reorganized the clusters based on their within-cluster mean BOLD signal.

### Modified reinstatement analysis

We performed a “modified” version of the standard reinstatement analysis, wherein movie event patterns were recalculated using “cluster patterns” derived from *k*-means clustering. For each time point in the group-average movie-viewing data, the original multi-voxel pattern at was replaced with the mean pattern of its assigned cluster (i.e., the cluster pattern). For example, if a time point was assigned to cluster 4 by the *k*-means clustering algorithm, the activity pattern at that time point was replaced with the mean pattern of cluster 4. A correlation matrix was then constructed for each subject by computing the correlation between a pair of recalculated movie event patterns and recall event patterns. These subject-level matrices were averaged to generate a group-average correlation matrix. In the group matrix, reinstatement strength was quantified as the average correlation of matching movie and recall event pairs (i.e., the diagonal of the matrix). This procedure was repeated for *k*-values ranging from 2 to 20, with *k*-means clustering performed anew for each of 10 iterations. The average correlation of matching event pairs for each *k*-value was averaged across 10 iterations. This value was then compared to 95% of the reinstatement strength from the original group matrix (i.e., “standard” reinstatement effects). The original group matrix was derived by computing each subject’s movie-recall event correlations using the original group-average movie-viewing data. This analysis was conducted separately for the left and right PMC, early auditory cortex, and early visual cortex ROIs.

### Identification of neural states: Recall

We performed *k*-means clustering on each individual’s recall data and compared the resulting recall cluster patterns to those obtained from the group-average movie-viewing data. For four subjects who recalled the movie scenes across two scanning runs, the data were concatenated across runs prior to analysis. Similar to the procedures previously used for identifying neural states during movie-viewing, the BOLD signal, averaged across vertices, was removed at each time point before clustering. For each subject, we aimed to identify the recall cluster labels that best matched the labels of movie cluster patterns derived from the group-average movie-viewing data. Recall cluster labels for each subject were reordered to maximize the average diagonal value of that subject’s movie cluster-by-recall cluster correlation matrix while keeping the movie cluster patterns in their original order. Subsequently, the average diagonal value was calculated from a group-level matrix, created by averaging across individuals’ matrices. Statistical significance for this value was tested using a permutation test, in which the recall cluster order for each subject was randomly shuffled before generating a new group-average matrix. The shuffling was performed after determining the recall cluster order that best matched the movie clusters. These procedures were repeated 10,000 times, constructing a null distribution for the average diagonal value of the group-average matrix from these permutations.

### Actions and social-affective features in movies

#### Stimuli and Procedures

314 online participants recruited from Prolific identified actions in a given 3-sec movie clip, and rated its social-affective properties. To create the clips, 3-second segments were extracted from the original movies at every 1.5 seconds (1 TR), resulting in a total of 1,848 video clips without audio. The clips were then divided into 77 subsets, each containing 24 different clips that were randomly selected from the pool of 1,848 clips. During the online task, each participant was randomly assigned one subset. Within the subset, the clips were presented in a randomized order. Participants were allowed to play a clip as many times as they wished.

#### Action labeling

For each clip, participants were asked to judge whether any character was present, including non-human animated characters or body parts, such as a hand or foot. This question was used (after data collection) to exclude participants who were not attending sufficiently. Next, participants were asked to list up to three actions that they observed in the clip (e.g., “walking”, “observing”, “searching”).

#### Social-affective feature labeling

We chose three social-affective features (sociality, valence, and arousal) based on prior findings that these features of everyday actions observed in naturalistic videos are critical for explaining the behavior-based dissimilarity structure of actions. Additionally, social-affective features are known to be processed at later stages during cortical information processing^49^. Participants provided social-affective ratings on a five-point scale in response to the following questions: “How social are the events in the video? (sociality; 1: not at all, 5: very social),” “How pleasant are the events in the video? (valence; 1: very unpleasant, 5: very pleasant),” “How intense are the events in the video? (arousal; 1: very calm, 5: very intense).” The order of the rating questions was randomized across participants.

#### Data cleaning

We excluded data from six participants whose accuracy on the question about the presence of a character was lower than 90%, retaining data from 309 participants. For the action label data, we excluded responses that identified an object (e.g., “camera”, “books”), a place (e.g., “office”), or characters (e.g., “a panel of contestants”, “lady in the car”) instead of an action. Additionally, we removed responses reflecting participants’ subjective judgments of the clip (e.g., “uniformity”, “stillness”, “kindness”) or emotions (e.g., “happy”, “looking concerned”). Adjectives describing specific facial expressions were excluded, while action verbs entailing facial expressions were retained. For example, the label “smile” was considered a valid action verb (e.g., people smile when they are happy), whereas the label “happy” was removed. When possible, we converted noun-based responses into action verbs; for instance, “discussion” was converted to “discuss.”

In the valence data, one participant assigned the value 3 (neutral) to all of the given 24 clips. Since no other problems were apparent in that participant’s responses to other questions, we excluded only the valence ratings for that participant.

### Voxel-wise encoding model estimation and validation

Three encoding models based on multiple linear regression were constructed: 1) an “action” model, 2) a “social-affective” model, and 3) a full model. During model estimation, each model was trained to predict PMC neural states (i.e., cluster patterns derived from group-average movie-viewing data) using actions, social-affective features, or both, separately for each hemisphere. For the “action” model, predictors were 50-dimensional GloVe^52^ word embedding vectors derived from action labels collected from online participants. For each time point (TR), embedding vectors of all action labels provided by participants for the two consecutive clips were averaged. Each time point was covered by two consecutive clips, as 3-sec clips were generated every 1.5 sec (1 TR) with a 1.5-sec overlap between consecutive clips during the data collection of actions and social-affective ratings. Repeated labels were included when generating a single embedding vector for actions at each time point. For example, if two participants provided the label “walking,” the embedding vectors for both instances were included in the average, giving greater weight to actions reported by more participants. For the “social-affective” model, predictors were ratings of sociality, valence, and arousal. These ratings were *z*-scored within each participant and then averaged across participants who watched the clips corresponding to that time point. Similar to the action model, each time point was covered by two successive 3-sec clips, resulting in a three-dimensional vector capturing the three social-affective features.

A leave-one-TR-out cross-validation scheme was adopted to test each model’s prediction performance using a total of 1,780 TRs (i.e., 1,780 iterations), after excluding periods with no action labels provided by any participant, including movie title scenes. Each model’s performance was evaluated by determining whether the predicted multi-voxel pattern for a held-out TR was most strongly correlated with its true cluster pattern label compared to all other possible cluster patterns, which were newly generated for each iteration. To avoid potential autocorrelation issues, during each iteration, data from 5 TRs on either side of the held-out TR was excluded from the model estimation and cluster pattern calculations. Cluster patterns were recalculated for each iteration, excluding the held-out TR and the surrounding 5 TRs.

## Data availability

The action labels and the social-affective ratings data will be made available upon publication.

## Code availability

Analysis scripts are available upon request to the corresponding author (Y.L.).

## Acknowledgments

We thank Drs. Christopher Honey and Leyla Isik for valuable intellectual input on data analyses and for discussions, and Euikwang Kim for assistance with collecting action labels and social-affective features from online participants. This work was supported by R01 MH133732 and MURI N00014-23-1-2086 awarded to J.C. and the William Orr Dingwall Dissertation Fellowship in the Foundation of Language awarded to Y.L.

## Author contributions

Y.L. and J.C. conceived and designed the study. Y.L. collected ratings data, analyzed the fMRI and ratings data, drafted the paper. H.L. preprocessed the fMRI data. Y.L., H.L., and J.C. edited the paper. J.C. supervised the project.

## Supplementary Information

**Supplementary Table 1.** ROI definition.

| Region of interest | Hemisphere | Schaefer parcel ID | Schaefer parcel name |
| --- | --- | --- | --- |
| Posterior medial cortex | Left | 154 | 17Networks_LH_DefaultA_pCunPCC_1 |
|  |  | 155 | 17Networks_LH_DefaultA_pCunPCC_2 |
|  |  | 156 | 17Networks_LH_DefaultA_pCunPCC_3 |
|  |  | 157 | 17Networks_LH_DefaultA_pCunPCC_4 |
|  |  | 158 | 17Networks_LH_DefaultA_pCunPCC_5 |
|  |  | 159 | 17Networks_LH_DefaultA_pCunPCC_6 |
|  |  | 160 | 17Networks_LH_DefaultA_pCunPCC_7 |
|  | Right | 363 | 17Networks_RH_DefaultA_pCunPCC_1 |
|  |  | 364 | 17Networks_RH_DefaultA_pCunPCC_2 |
|  |  | 365 | 17Networks_RH_DefaultA_pCunPCC_3 |
|  |  | 366 | 17Networks_RH_DefaultA_pCunPCC_4 |
|  |  | 367 | 17Networks_RH_DefaultA_pCunPCC_5 |
| Early visual cortex | Left | 7 | 17Networks_LH_VisCent_Striate_1 |
|  |  | 18 | 17Networks_LH_VisPeri_StriCal_1 |
|  |  | 19 | 17Networks_LH_VisPeri_StriCal_2 |
|  |  | 20 | 17Networks_LH_VisPeri_ExStrSup_1 |
|  | Right | 207 | 17Networks_RH_VisCent_Striate_1 |
|  |  | 218 | 17Networks_RH_VisPeri_StriCal_1 |
|  |  | 219 | 17Networks_RH_VisPeri_StriCal_2 |
| Early auditory cortex | Left | 44 | 17Networks_LH_SomMotB_Aud_1 |
|  |  | 45 | 17Networks_LH_SomMotB_Aud_2 |
|  |  | 46 | 17Networks_LH_SomMotB_Ins_1 |
|  | Right | 244 | 17Networks_RH_SomMotB_Aud_1 |
|  |  | 245 | 17Networks_RH_SomMotB_Aud_2 |
|  |  | 246 | 17Networks_RH_SomMotB_Ins_1 |
| Angular gyrus | Left | 149 | 17Networks_LH_DefaultA_IPL_1 |
|  |  | 150 | 17Networks_LH_DefaultA_IPL_2 |
|  |  | 173 | 17Networks_LH_DefaultB_IPL_1 |
|  |  | 174 | 17Networks_LH_DefaultB_IPL_2 |
|  |  | 188 | 17Networks_LH_DefaultC_IPL_1 |
|  | Right | 359 | 17Networks_RH_DefaultA_IPL_1 |
|  |  | 360 | 17Networks_RH_DefaultA_IPL_2 |
|  |  | 385 | 17Networks_RH_DefaultC_IPL_1 |
|  |  | 386 | 17Networks_RH_DefaultC_IPL_2 |

**Supplementary Figure 1.**
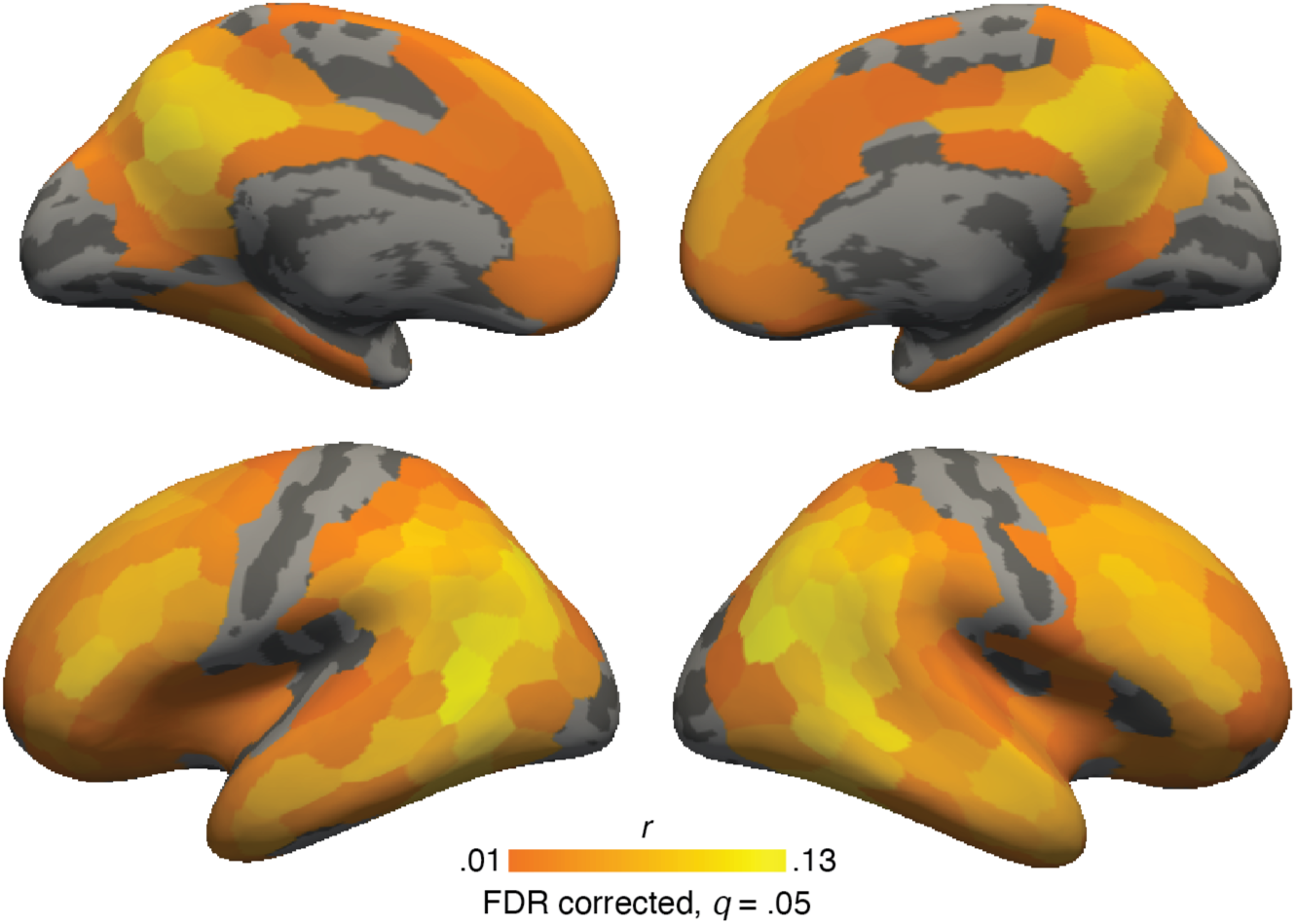
Parcel-wise within-subject standard memory reinstatement analysis results. We performed an analysis in which the correlation for the matching pairs of movie and recall event patterns, averaged across events (i.e., reinstatement strength), was calculated for each cortical parcel using each subject’s own data. The surface shows the reinstatement strength averaged across subjects at the parcel level after FDR correction at *q* = .05. Statistical significance was evaluated using a randomization test (shuffling of movie event labels; see *Whole-brain within-subject standard reinstatement analysis* in Methods). Robust reinstatement strength was observed in core DMN areas, such as in bilateral PMC, which is a key focus of our study, and the bilateral angular gyrus.

**Supplementary Figure 2.**
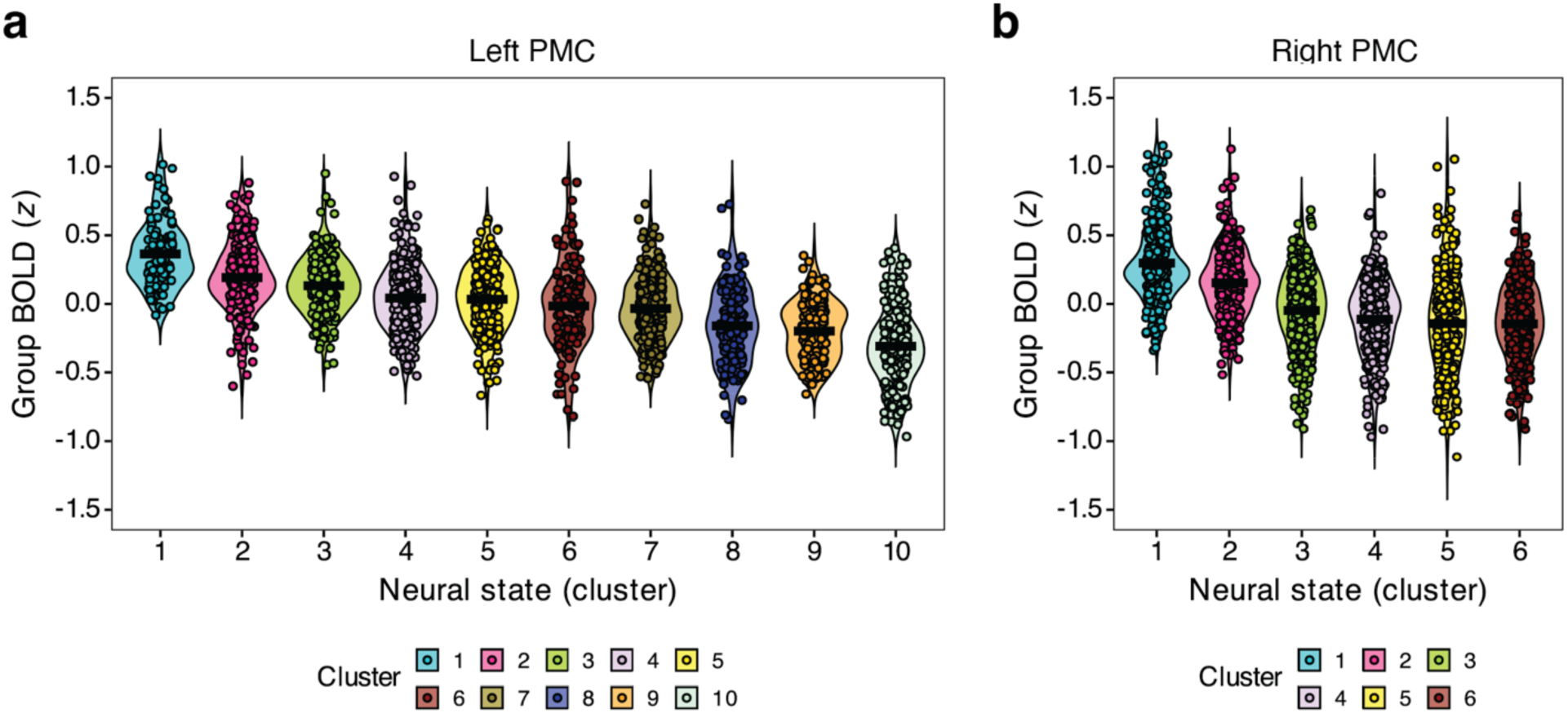
Clusters organized by BOLD signal levels. The distribution of BOLD signals at individual time points is shown for each cluster (i.e., neural state). Since the *k*-means clustering algorithm randomly assigns cluster numbers, we reorganized the clusters (without altering membership) to create a numbering scheme based on the average BOLD signal within each cluster. Specifically, *cluster 1* corresponds to the cluster with the highest BOLD signal, while *cluster k* corresponds to the cluster with the lowest BOLD signal. The term “BOLD signal” refers to the group BOLD signal, which is the average activity across vertices in the group-averaged movie-viewing vertex-by-time data matrix, used for *k*-means clustering in the prior neural state identification step. **a.** The BOLD signal distribution of the left PMC data, with results for *k* = 10 for illustrative purposes. Each violin plot includes dots showing data from individual time points. **b.** The results for *k* = 6 for the right PMC data.

**Supplementary Figure 3.**
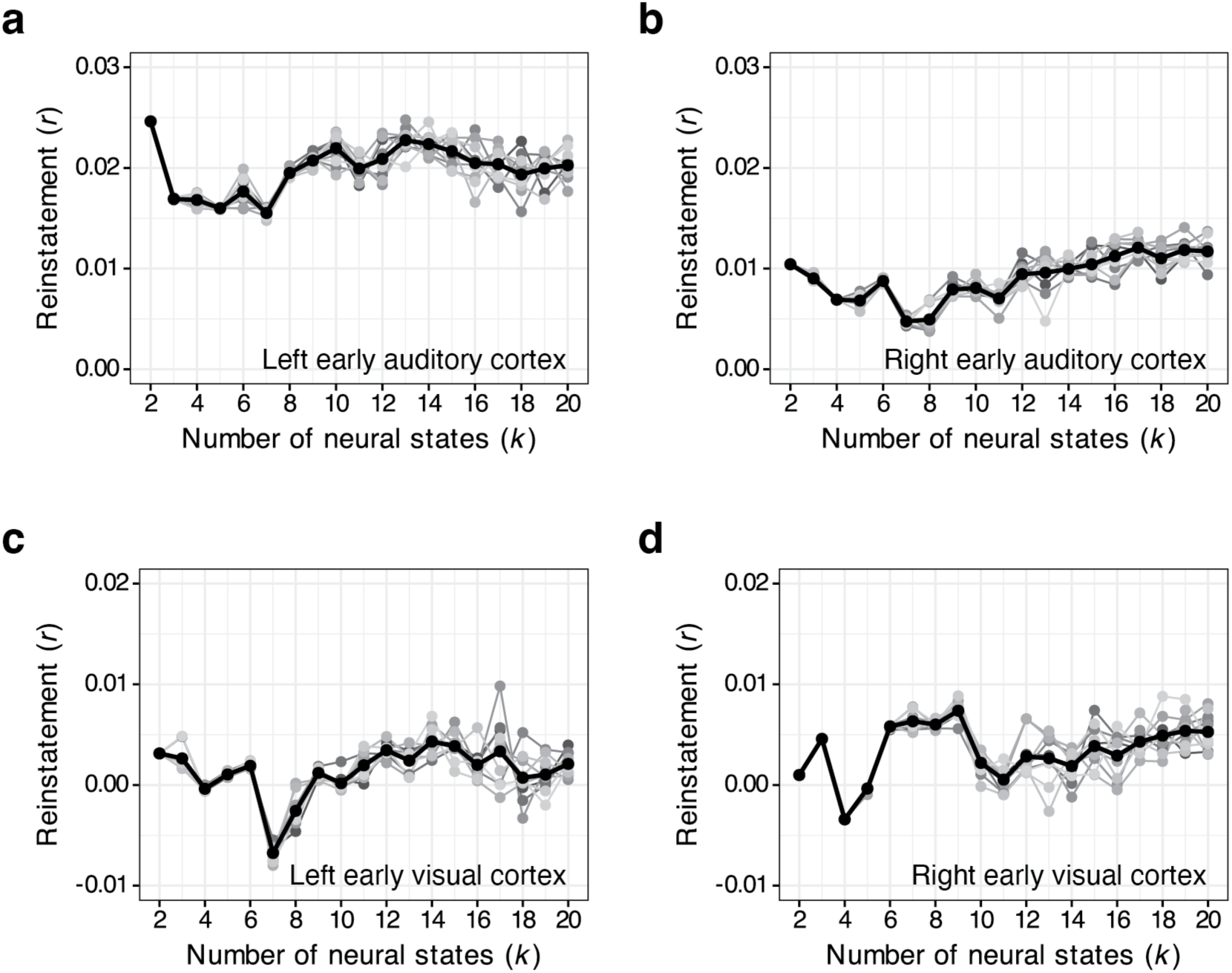
Modified reinstatement analysis results for sensory processing areas. In each plot, the black bold line indicates the results averaged across 10 iterations. **a-b.** Early auditory cortex. **c-d.** Early visual cortex. These regions did not show strong reactivation in either the within-subject standard analysis (Supplementary Fig. 1) or the modified analysis, as shown on the y-axis of the current figure.

**Supplementary Figure 4.**
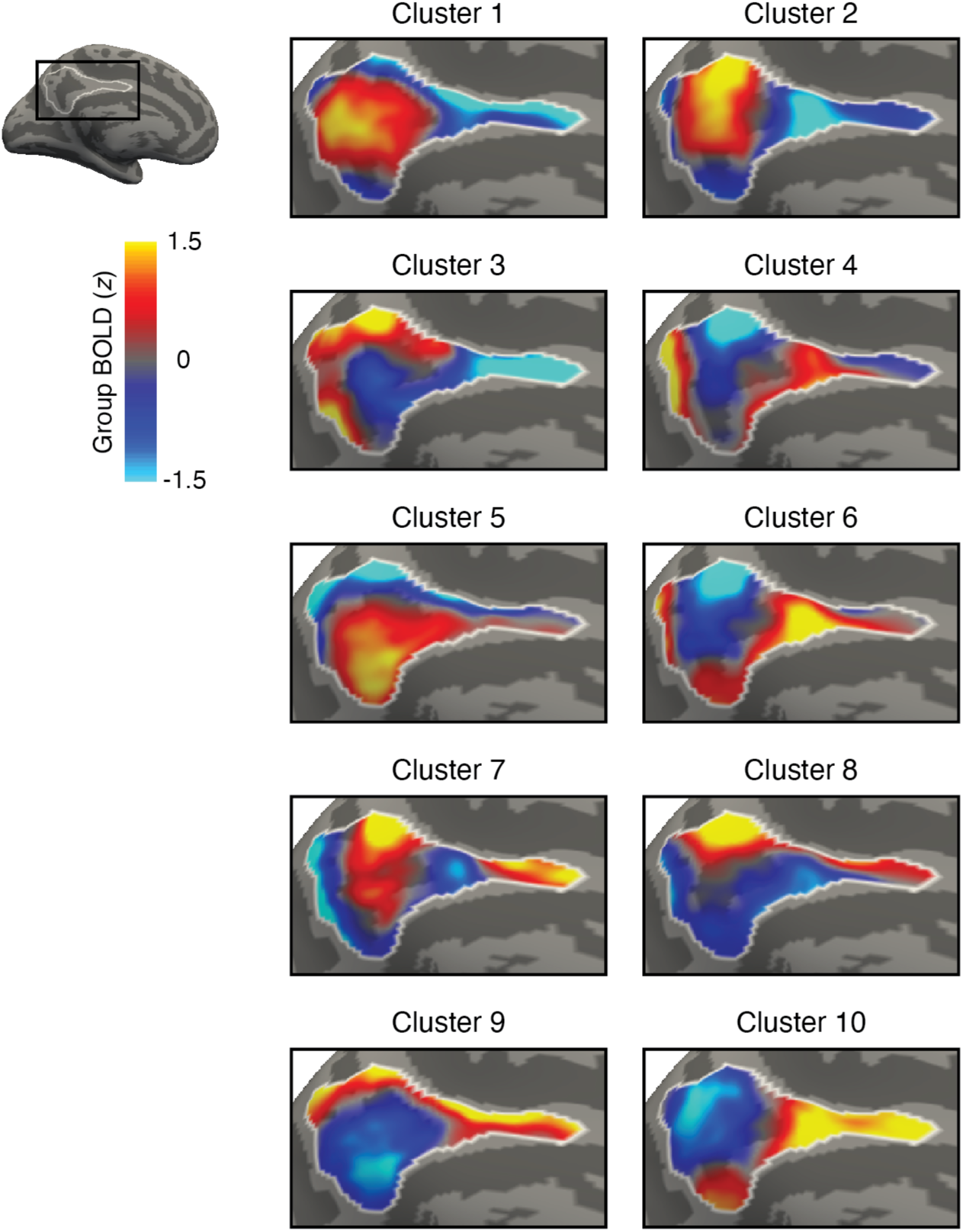
Cluster patterns identified in the left PMC when *k* = 10. These cluster patterns were generated from a single iteration of *k*-means clustering for visualization purposes. The same *k*-means clustering solution was used to create other figures related to the left PMC data.

**Supplementary Figure 5.**
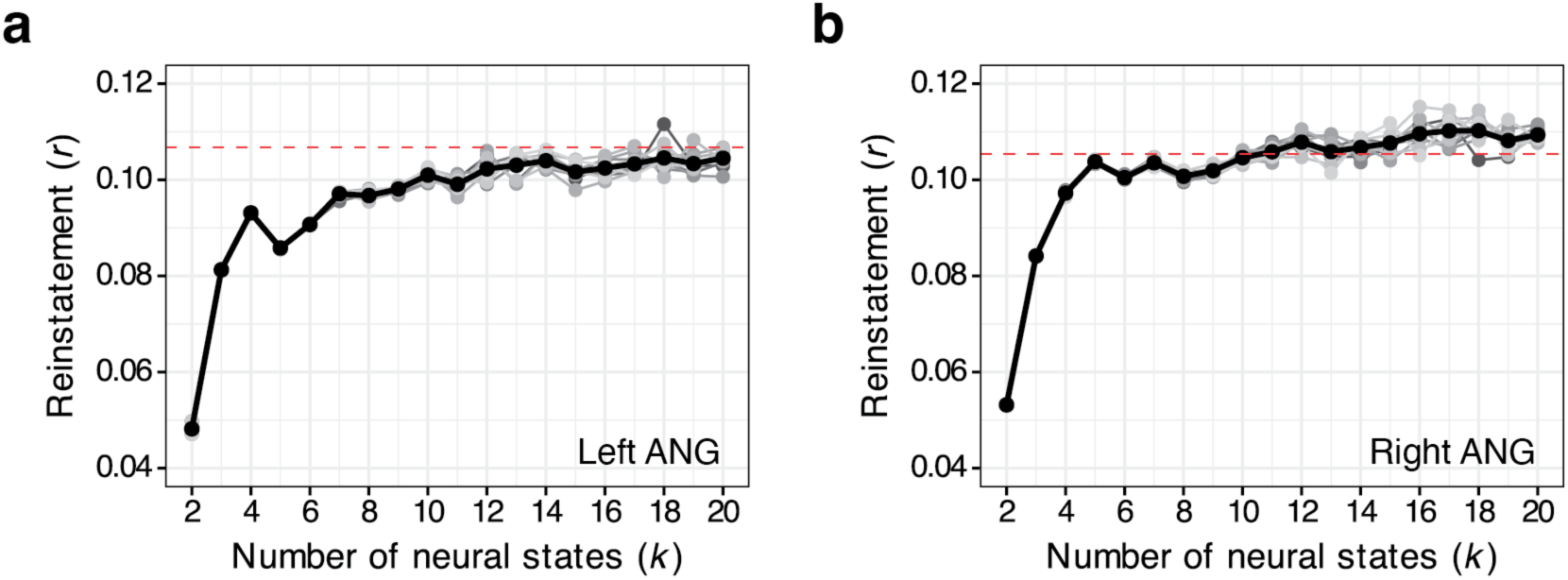
Modified reinstatement analysis results for the angular gyrus (ANG). In each plot, the red dashed line indicates 95% of the standard reinstatement effects for each area, while the black bold line indicates results averaged across 10 iterations. **a** In the left ANG, the reinstatement effects were not reliably achieved over 10 iterations, even with 20 neural states. **b** In the right ANG, 11 neural states were required to observe 95% of its standard effects. Interestingly, we observed lateralization: a smaller number of neural states were needed in the right hemisphere to observe the effects, consistently across both the right PMC and the right ANG.

**Supplementary Figure 6.**
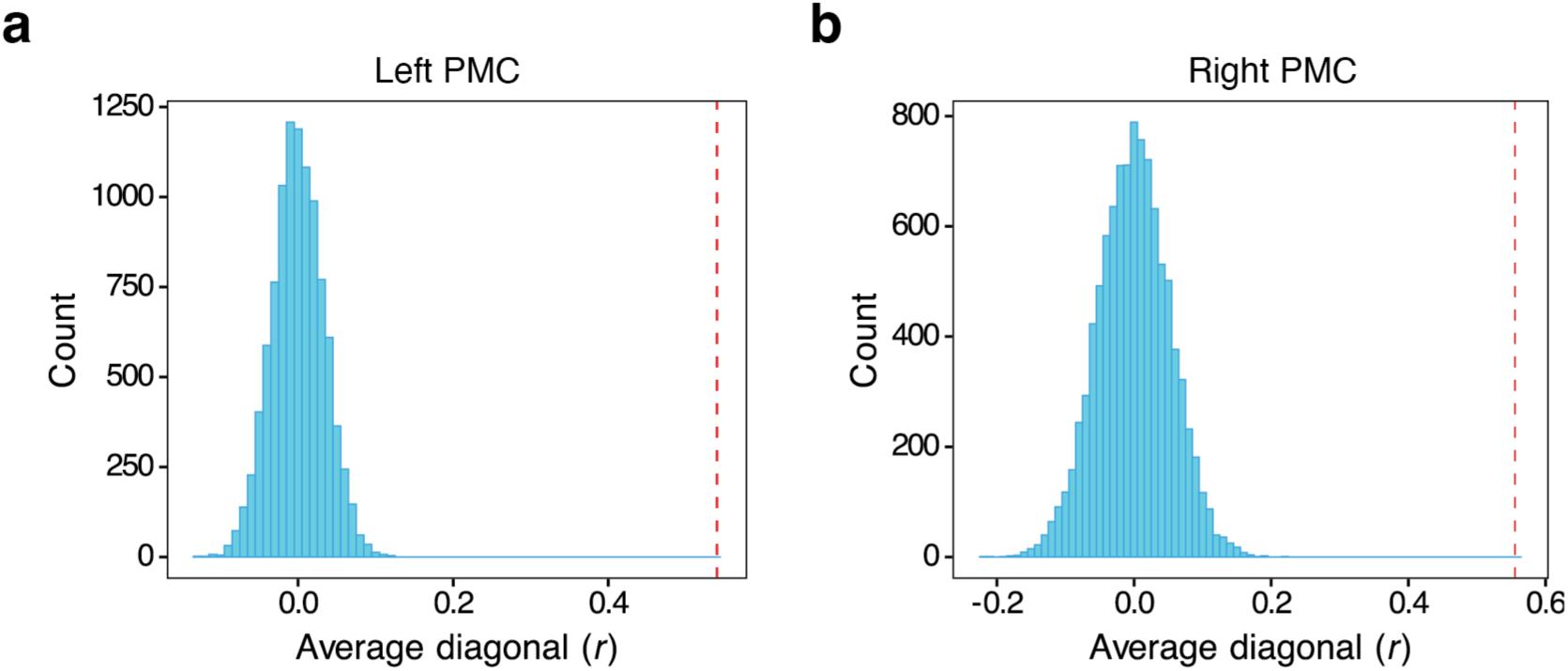
Null distribution constructed from a permutation test. In each plot, the red vertical line indicates the observed average diagonal value of the group-average matrix after the best match procedure. In contrast, the distribution of permuted data is centered around zero. **a** Left PMC. **b** Right PMC.

**Supplementary Figure 7.**
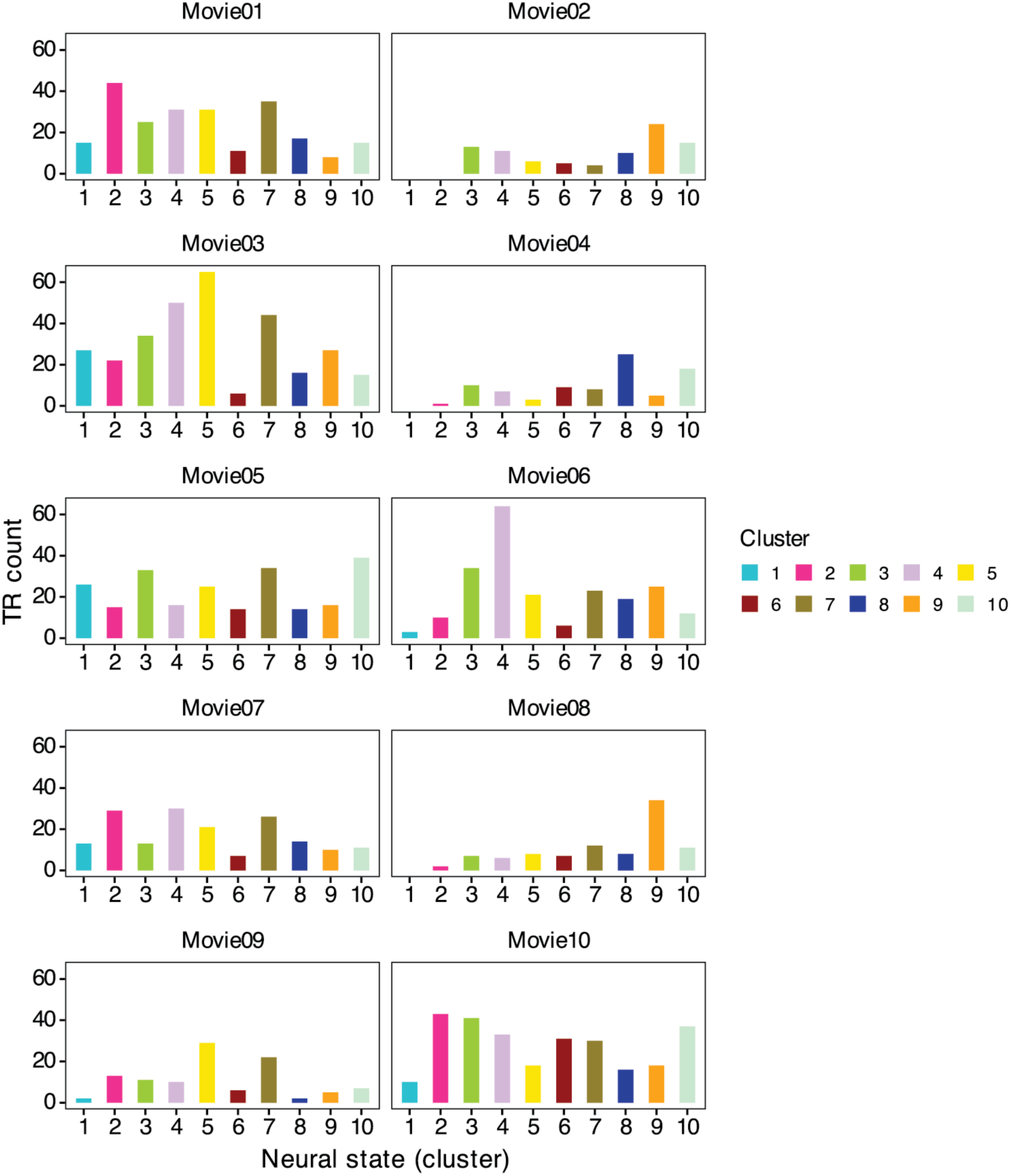
Cluster occurrences in each movie for the left PMC when *k* = 10. Each bar shows the number of time points (TRs) assigned to each cluster. Time points from the title scene periods were excluded when calculating neural state occurrences.

**Supplementary Figure 8.**
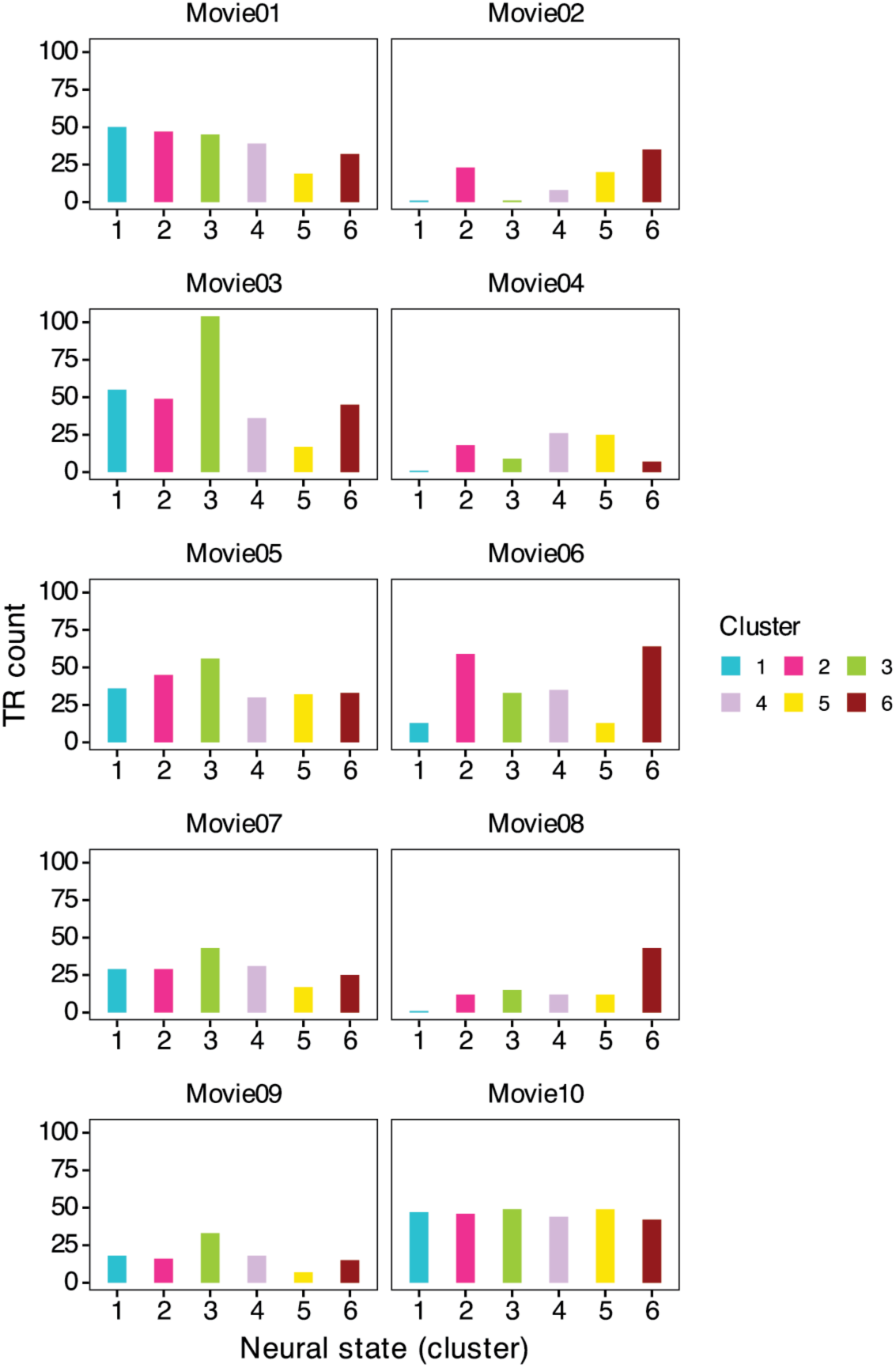
Cluster occurrences in each movie for the right PMC when *k* = 6. Each bar shows the number of time points (TRs) assigned to each cluster. Time points from the title scene periods were excluded when calculating neural state occurrences.

**Supplementary Figure 9.**
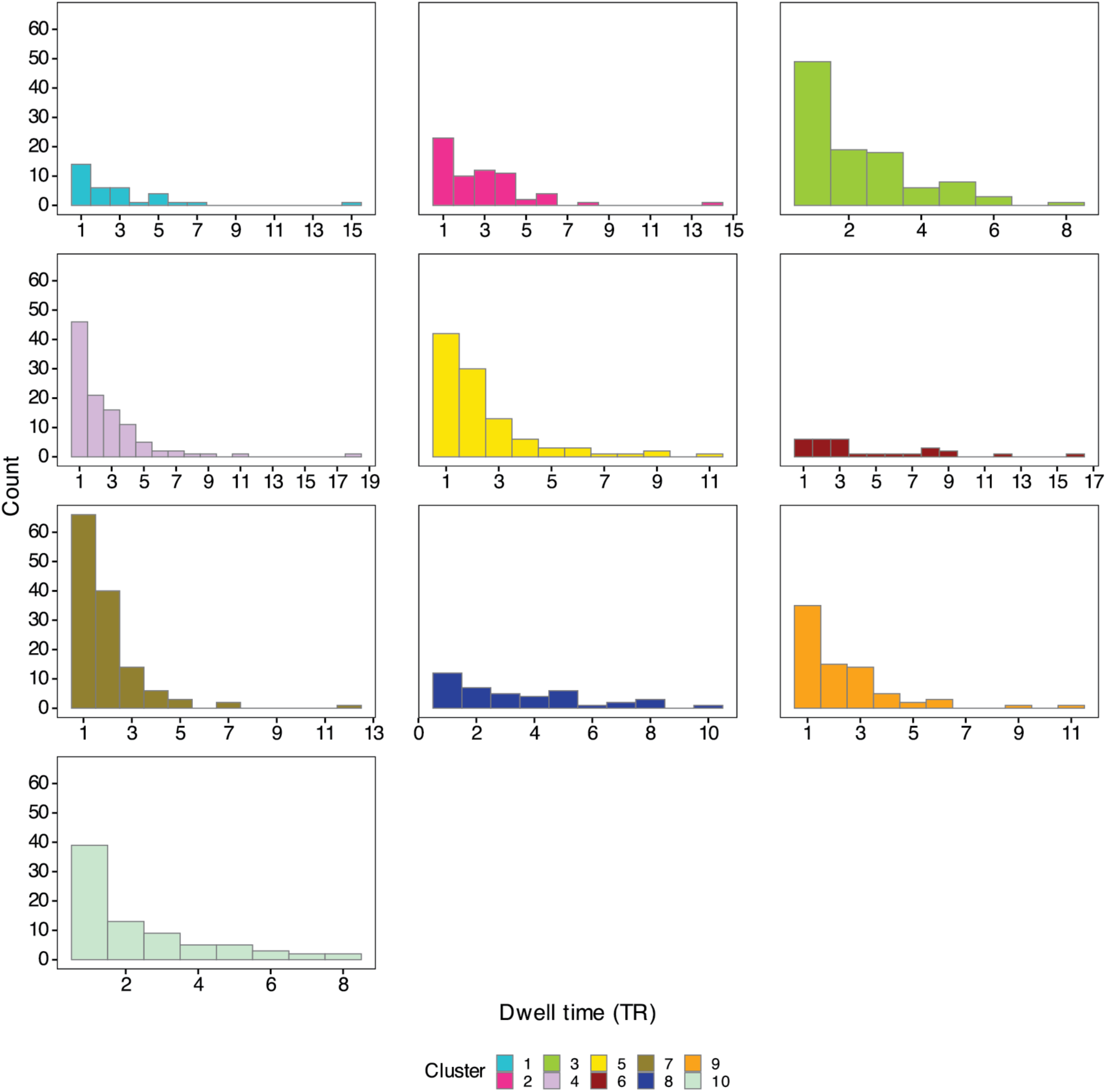
The distribution of dwell times for each neural state in the left PMC. Each graph shows the histogram of dwell times for each neural state (cluster), distributed across all 10 movies, with the chosen value of *k* = 10. The y-axis shows the number of cluster segments for each corresponding bin. The neural states were predominantly transient.

**Supplementary Figure 10.**
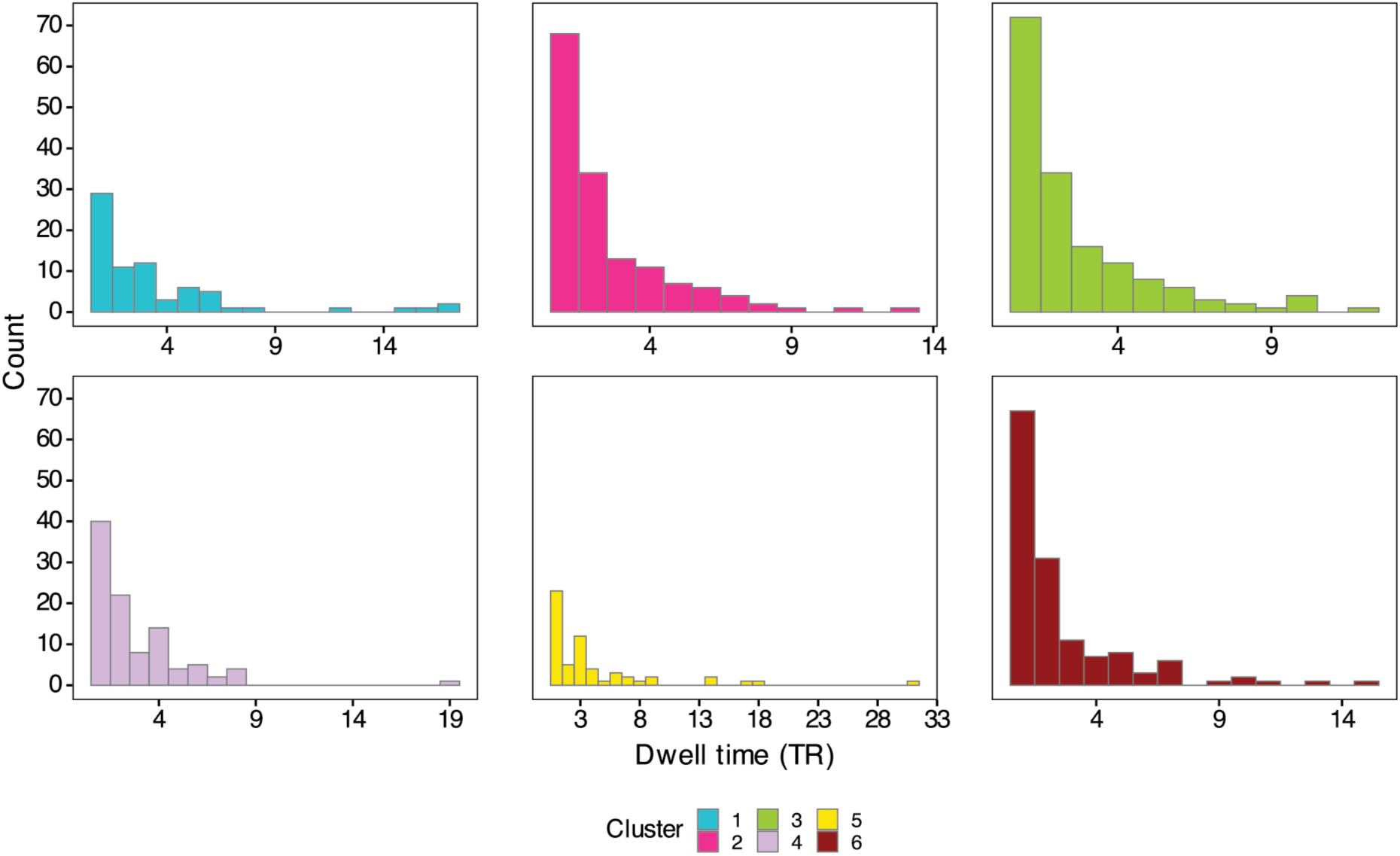
The distribution of dwell times for each neural state in the right PMC. Each graph shows the histogram of dwell times for each neural state (cluster), distributed across all 10 movies, with the chosen value of *k* = 6. The y-axis shows the count of cluster segments in each bin. Similar to the left PMC (Supplementary Fig. 9), the neural states showed transient characteristics.

**Supplementary Figure 11.**
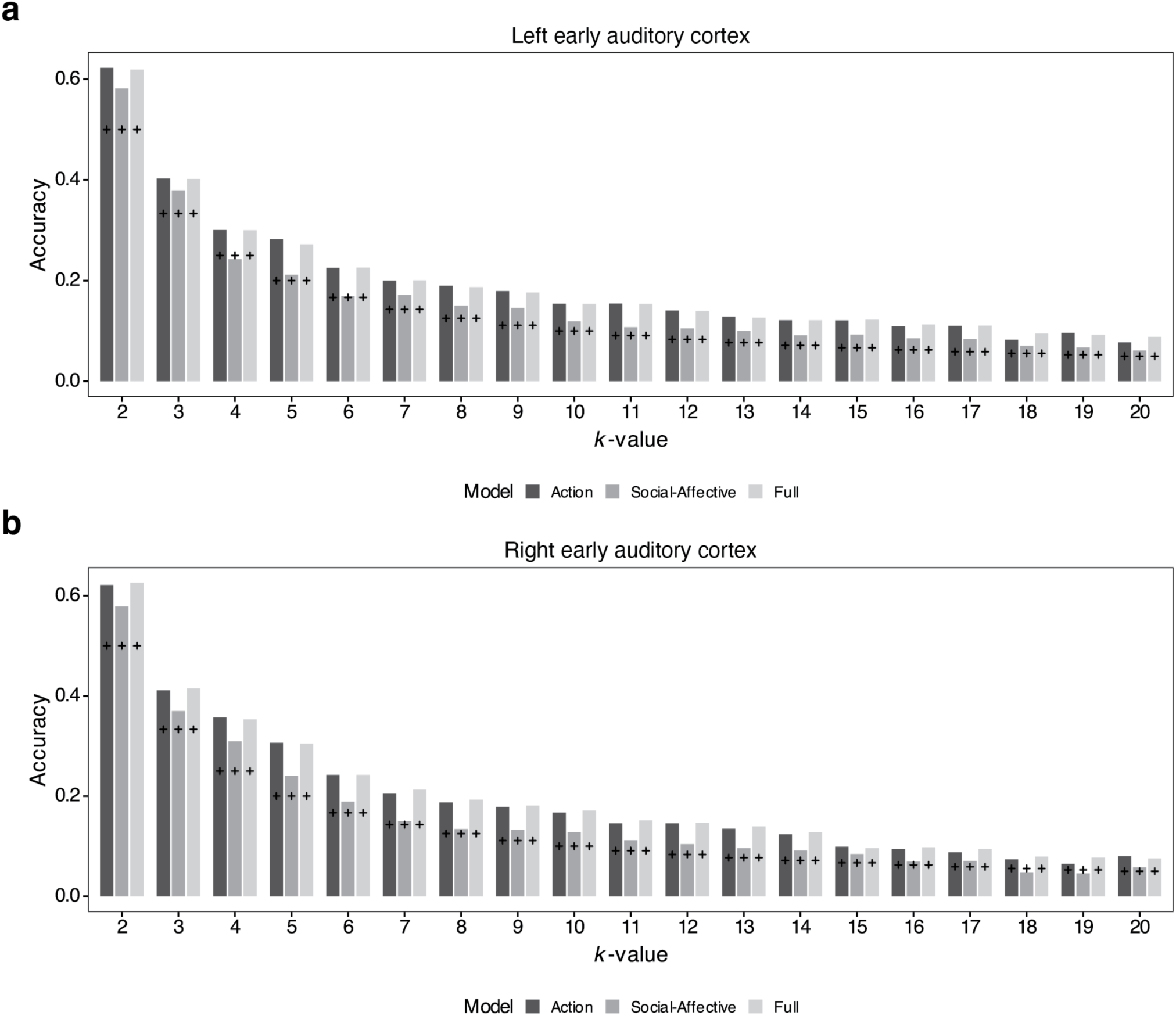
Encoding model analysis results for early auditory cortex. Results are shown for all *k*-values since these regions did not show strong memory reactivation, and no specific *k*-value (i.e., the number of neural states) was selected for these ROIs from the modified version of the reinstatement analysis. We observed that actions and social-affective features in the movies were associated with neural states for most *k*-values in these regions, as well as the left and right PMC, despite their relatively low reinstatement effects. The crossbar marker indicates the chance level for each *k*-value.

**Supplementary Figure 12.**
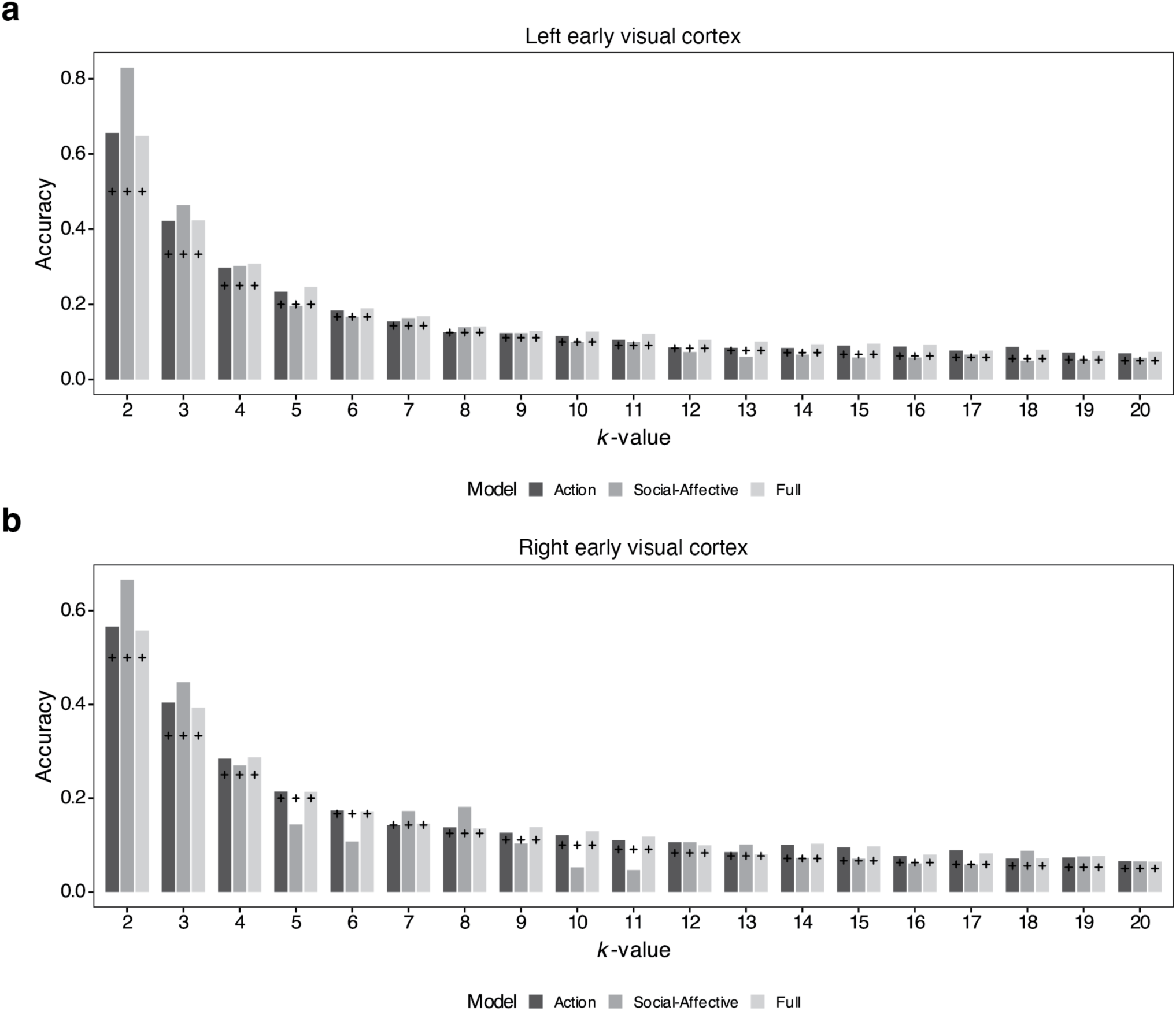
Encoding model analysis results for early visual cortex. Similar to the early auditory cortex (Supplementary Fig. 11), the results are presented for all *k*-values because weak memory reactivation was observed in these regions, and no specific *k*-value was chosen for these ROIs in the modified reinstatement analysis. The crossbar marker indicates the chance level for each *k*-value.

**Supplementary Figure 13.**
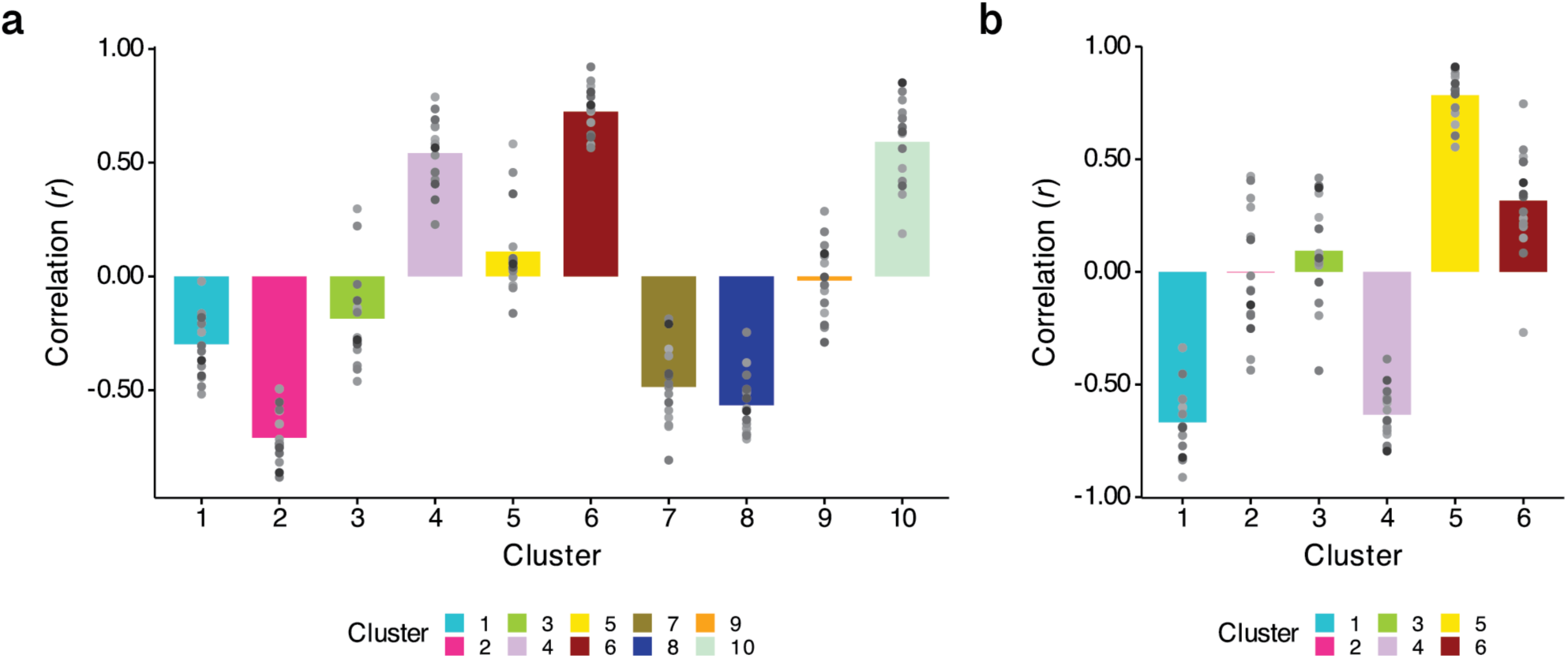
Correlation between each subject’s movie cluster pattern and their own recall boundary pattern. We observed one cluster pattern that was most strongly correlated with a recall boundary pattern when individual movie-viewing data, rather than group-average data, were used to calculate the movie cluster patterns. Group-derived cluster labels were applied to each subject’s movie-viewing data for these calculations. **a** Cluster 6 showed the strongest correlation in the left PMC when *k* = 10. **b** Cluster 5 showed the strongest correlation in the right PMC when *k* = 6. Gray dots indicate individual subjects’ data points.

